# Structural and functional alterations associated with the LRRK2 G2019S mutation revealed in structured human neural networks

**DOI:** 10.1101/2020.05.02.073726

**Authors:** Vibeke Devold Valderhaug, Ola Huse Ramstad, Rosanne van de Wijdeven, Kristine Heiney, Stefano Nichele, Axel Sandvig, Ioanna Sandvig

## Abstract

Mutations in the LRRK2 gene have been widely linked to Parkinson’s disease. The G2019S variant has been shown to contribute uniquely to both familial and sporadic forms of the disease. LRRK2-related mutations have been extensively studied, yet the wide variety of cellular and network events directly or indirectly related to these mutations remain poorly understood. In this study, we structured multi-nodal human neural networks carrying the G2019S mutation using custom-designed microfluidic chips coupled to microelectrode-arrays. By applying live imaging approaches, immunocytochemistry and computational modelling, we have revealed alterations in both the structure and function of the resulting neural networks when compared to controls. We provide first evidence of increased neuritic density associated with the G2019S LRRK2 mutation, while previous studies have found either a strong decrease, or no change, compared to controls. Additionally, we corroborate previous findings regarding increased baseline network activity compared to control neural networks. Furthermore, we can reveal additional network alterations attributable to the specific mutation by selectively inducing transient overexcitation to confined parts of the structured multi-nodal networks. These alterations, which we were able to capture both at the micro- and mesoscale manifested as differences in relative network activity and correlation, as well as in mitochondria activation, neuritic remodelling, and synaptic alterations. Our study thus provides important new insights into early signs of neural network pathology significantly expanding upon the current knowledge relating to the G2019S Parkinson’s disease mutation.

## 1. Introduction

Mutations in the leucine-rich repeat kinase 2 (LRRK2) (PARK8 locus) gene are linked to both late-onset familial and sporadic forms of Parkinson’s disease (PD) (Healy et al., 2008). Although rare, LRRK2 gene mutations have been termed a potential “Rosetta stone” of parkinsonian disorders as all of the major pathologies related to parkinsonism have been observed, in addition to there being end-stage variability, within families carrying the same pathogenic variant (Trinh et al., 2006). Moreover, the Lewy body and Lewy neurite pathology commonly observed in brain autopsies from PD patients are rarely observed in relation to LRRK2 mutations (Moore, 2008). The LRRK2 gene is expressed both in the brain and in other tissues throughout the body and is translated into the LRRK2 protein, which has enzymatic kinase activity involved in a range of cellular processes (Trinh et al., 2006). Pathogenic variants are postulated to augment this kinase activity, resulting in a toxic gain-of-function through an increase in both autophosphorylation and phosphorylation of LRRK2 substrates (Trinh et al., 2006, Sheehan and Yue, 2018, Jaleel et al., 2007, Zhao et al., 2012, Smith et al., 2006, Greggio et al., 2006, Steger et al., 2016).

Studies using both *in vitro* and *in vivo* models of PD suggest that synaptic alterations and axonal dysfunction represent the earliest detectable signs of the disease (MacLeod et al., 2006, Cheng et al., 2010, Dagda et al., 2014) and that initiation of pathology at the axon terminals might signify the start of the retrograde degeneration of the neurons (Tagliaferro and Burke, 2016, Sheehan and Yue, 2018). This also fits with the latest estimates compiling observations from several independent studies, concluding with 50-70% loss of striatal terminals at symptom onset of PD, while “only” 30% of dopaminergic neurons in the substantia nigra are lost at the same timepoint (as opposed to the often reported 60-80% dopaminergic neuron loss) (Cheng et al., 2010). Moreover, there is now abundant evidence that the molecular mechanisms of axonal degeneration are distinct from those of programmed cell death, implying that the two mechanisms should be considered separately, both in disease modelling and therapeutics (Cheng et al., 2010, Tagliaferro and Burke, 2016). Dendritic spine loss and shortening and simplification of the dendritic arbor are also regularly observed in post mortem tissues from patients with Alzheimer’s disease and amyotrophic lateral sclerosis (Brizzee, 1987, Stephens et al., 2005, Baloyannis, 2006, Sasaki and Iwata, 2007, Cherra et al., 2013, Dagda et al., 2014, Genc et al., 2017). Such changes are often accompanied by a loss or impairment of dendritic mitochondria, a feature which is central in PD pathogenesis (Bose and Beal, 2016, Verma et al., 2017, Singh et al., 2019). Furthermore, these alterations have been linked to increased excitatory stimulation and calcium handling (Caudle and Zhang, 2009, Verma et al., 2018).

The advancement and availability of tools for morphogenetic neuroengineering now enable modelling of selected pathological aspects of neurodegenerative disease in human neural networks in vitro. In this study, we utilize healthy human cortical neural networks and equivalent networks carrying the Parkinson’s related LRRK2 G2019S mutation to study early expression of pathology, both at the micro- and mesoscale. Among the LRRK2 mutations, the particular G2019S mutation represents the most commonly identified cause of late-onset PD, and has been shown to contribute uniquely to both familial and sporadic forms of the disease (Kachergus et al., 2005, Bouhouche et al., 2017, Trinh et al., 2006, Moore, 2008, Healy et al., 2008, Gilks et al., 2005, Lesage et al., 2005, Ozelius et al., 2006, Di Fonzo et al., 2005, Nichols et al., 2005). Moreover, it has been shown to result in differential regulation of a variety of cellular pathways highly relatable to several aspects of PD pathology, among which are axonal guidance, cytoskeletal transport, cell growth, differentiation and communication (Habig et al., 2013, Habig et al., 2008).

In this study we investigate whether in vitro neural networks carrying the LRRK2 G2019S mutation display a different structural and functional profile compared to healthy controls during development and in response to induced perturbation consistent with excitotoxicity. To address these aims, we structured human neural networks with and without the LRRK2 G2019S mutation using tailor-made multi-nodal microfluidic chips (van de Wijdeven et al., 2018, van de Wijdeven et al., 2019). These microfluidic chips incorporate axon tunnels and synaptic compartments interconnecting three cell chambers, thus allowing selective manipulation of the neural network. Furthermore, each microfluidic chip incorporates a microelectrode-array (MEA), which enables electrophysiological recording of neural network activity within different interconnected nodes. We have subsequently monitored early network structure, function and dysfunction related to the G2019S LRRK2 mutation using this platform. Importantly, based on the multiple-hit hypothesis of PD (Carvey et al., 2006, Sulzer, 2007, Patrick et al., 2019), according to which a combination of genetic susceptibility and environmental factors may contribute to the onset and progression of the disease, we have selectively induced a transient, topologically confined neural overexcitation event, monitored network responses and revealed associated alterations at the micro- and mesoscale.

## 2. Materials and Methods

### Structuring cortical neural networks using microfluidics chips with directional inter-nodal connectivity

Control human induced pluripotent stem cell (iPSC)-derived H9N neural stem cells (NSCs) (ax0019) and iPSC derived H9N NSCs homozygously carrying the LRRK2 G2019S (GGC>AGC) mutation (ax0310) (Axol Bioscience, Cambridge, United Kingdom) were cultured and expanded on 0.01 % poly-L-ornithine (PLO) (Sigma) and L-15 laminin (L15 medium containing 1:60 natural mouse laminin and 1:41 sodium bicarbonate) coated culture vessels in neural expansion medium (ax0030) supplemented with human FGF2 and EGF (ax0047 and ax0047X), and kept in a standard humidified air incubator (5% CO^2^, 20%O^2^, 37°C) (full cell culture protocol, as well as further information on each cell line available in the supplementary data). Each microfluidic chip was coated with the same combination of PLO and L-15 laminin and seeded with 1.1×10^5^ NSCs (37000 cells per cell chamber), from which point synchronous differentiation and maturation of the NSCs into cortical neurons was carried out until day 15, using an NSC reagent bundle and media (ax0101) in accordance with the manufacturer’s protocol.

### Excitatory stimulation of structured neural networks using Kainic Acid

Fifteen days post seeding, the cortical networks in the microfluidics chips were stimulated with Kainic acid (KA) and subsequently investigated with live staining assays. KA (10μM) was applied to the top cell chamber for 30 minutes, after which all cell chambers were washed 3x with Dulbecco’s phosphate buffered saline (PBS) and resupplied with media. A flow barrier created by a 10μl media level difference between the stimulated chamber and the nonstimulated chambers ensured the confinement of the KA to the top cell chamber only. The same procedure was carried out for each cell line using only PBS as a control condition. A total reactive oxygen species (ROS) assay kit 520nm (Thermo Fisher Scientific) fluorescently labeling ROS production was applied to verify a cellular response and the confinement of the stimulation by fluorescence microscopy (EVOS FL auto 2, Invitrogen, California, United States), where the microscope was set to image simultaneously in each of the microfluidics chips chambers every 10 minutes for 1 hour immediately following KA stimulation.

### Axonal mitochondrial distribution in structured neural network

To investigate the mitochondria distribution in the control and LRRK2 neural networks, 0.1% Tetramethylrhodiamine (TMRM, T668, Invitrogen) was applied for 30 minutes at 37°C to all microfluidic chip chambers, rinsed in PBS, and imaged using a Zeiss 510 META Live confocal scanning laser microscope in a heated chamber (37°C). As a baseline measure, 3 image series were taken every 10 minutes, where an image was taken every second for 1 consecutive minute. Thirty minutes after a confined KA stimulation of the cortical neurons in the top chamber (as described in the previous section), the imaging procedure was repeated.

### Cell viability assay

On day 16 (i.e. 24 hours post KA stimulation), a LIVE/DEAD viability/cytotoxicity kit (MP03224, Invitrogen) was applied to the neural networks in the microfluidics chips to determine whether the KA stimulation was sublethal. 0.8ul Ethidium homodimer-1 (2mM in DMSO/H_2_O 1:4) and 0.4μl Calcein AM (4mM in anhydrous DMSO) was diluted in 2ml PBS and applied to all chambers in the microfluidic chips for 15 minutes in 37°C. The fluorescently labelled neural networks were then washed with PBS and imaged (EVOS FL auto 2).

### Immunocytochemistry of the structured neural networks

24 hours post KA stimulation (or PBS as a control condition), structured neural networks from both the control and LRRK2 group were fixed and used for immunocytochemistry assays to assess the neurite and spine morphology. For fixation, 2% paraformaldehyde (PFA) was applied for 15 minutes followed by 4% PFA for 10-minutes and 3×15 minute washes at room temperature (RT). Blocking solution consisting of PBS with 5% normal goat serum (NGS) and 0.3% Triton-X was applied for 2 hours at room temperature and was followed by overnight incubation in primary anti-body solution (PBS with 1% NGS, 0.1% Triton-X) in 4°C. The following antibodies were used: Rabbit anti-Piccolo antibody (1:400) (ab20664), mouse-anti PSD95 (1:200) (ab13552), rabbit anti-CaMKII (1:250) (ab134041), rabbit anti-GRIK5 (1:200) (PA-5-41401), rabbit anti-Glutamate receptor 1 (AMPA) (1:500) (ab109450), mouse anti-MAP2 (1:400) (131500, Thermo Fisher Scientific), mouse anti-beta III tubulin (1:400) (ab119100), rabbit anti-total alpha synuclein (ab131508), mouse anti-mitochondria (ab3298), and chicken anti-neurofilament heavy (1:1000) (ab4680). The structured neural networks were then washed 3×15 minutes in PBS, and incubated for 3 hours in secondary antibody solution (PBS with 1% NGS, 0.1% Triton-X) in the dark, at RT. A combination of Alexa Fluor™ 488, 568, 647 secondary antibodies (Thermo Fisher, MA, USA) were used at a dilution of 1:1000. CytoPainter Phalloidin 647 (1:500) (ab176759) was added for the final 20 minutes of incubation, and Hoechst (1:10000) was added for the final 5 minutes before another 3×15 min wash in PBS was conducted. To avoid dilution of the solutions used for these immunocytochemistry procedures by the media already present in the axon tunnels of the microfluidic chips, some of the appropriate solution was used to flush through the tunnels prior to the actual incubation with each relevant solution. Images used for separate quantification of the fluorescently immunolabeled neural networks were taken using either a Zeiss Axiovert 1A fluorescent microscope (Carl Zeiss, Germany) with a 100x/1.3 oil objective or a Zeiss (510 META Live) confocal laser scanning microscope with a 63X/1.4 oil objective. ImageJ, MatLab and PowerPoint were used to post-process the images.

### Image analysis

Analysis of ROS expression was done using the Fiji plugin Particle analyzer. The two-channel fluorescent images from the viability assay were merged, adjusted for brightness/contrast for maximal separation and clarity of both signals, and the cells manually counted using the Cell counter Fiji-plugin. The area close to the active zone in the microfluidic chip was selected for analysis as this area showed better separation of the signals due to a consistent lower cell density across all conditions. The number and size of the TMRM-labelled mitochondria within the axonal tunnels were extracted using a simple image analysis pipeline implemented in MATLAB. First, nonuniform background illumination was removed by applying a top hat filter. The image was then binarized by Otsu thresholding, and the 8-connected components were extracted from the resulting binary image. Artefacts at the edges of the images, which tended to be large and elongated, were removed by eliminating components if their size exceeded 5 μm^2^ or eccentricity exceeded 0.995. The number of mitochondria was then extracted as the number of remaining 8-connected components in the image. The size of each detected mitochondrion was computed from the number of pixels comprising each as-detected component. Additional analysis of fixed samples from this experiment were analyzed in Fiji using ROI manager. Two channel 100X images of fluorescent mitochondria and total alpha synuclein in the neural networks were binarized by Otsu thresholding, and the area covered by the fluorescence in each channel was selected and measured using the ROI manager. A ratio of the area covered by mitochondria/total alpha synuclein was calculated and used for statistical analysis in Prism8 (GraphPad, California, United States).

The fluorescence images of Piccolo-immunolabelled neurites were analyzed using a semiautomated process implemented in MATLAB and Fiji to count the number of neuritic boutons present in the images. Most boutons were counted in an automated fashion in MATLAB. Top hat filtering was applied to suppress nonuniform background illumination before the contrast of the resulting images was enhanced using adaptive histogram equalization. The contrast-enhanced images were binarized using Otsu thresholding, and salt noise in the binarized images was removed by median filtering. Neurite fragments were then joined by morphological closing, and any remaining small fragments were removed by applying hole filling to the inversion of the resulting image. Thinning was then applied to obtain a skeleton of the neurites in the image. From this thinned image, endpoint detection was used to obtain a preliminary bouton count. Because the automated analysis missed some boutons, the endpoint labelled images were visually inspected and the missed boutons were manually counted using the Cell counter in Fiji and added to the final results. The area covered by the neurites was calculated by means of the Particle analyzer after binarization with Otsu thresholding. Together, the boutons counted divided by the area covered with neurites created a ratio used for statistical analysis. For the measure of co-occurring Piccolo and PSD95 immunolabelling, an automatic threshold was applied for each channel (Otsu for Piccolo and Triangle for PSD95) in Fiji, the thresholded areas selected as ROIs, and areas containing both ROIs were selected for particle analysis. A cut-off at 15μm was set as an upper limit, and the number and average size measurement from each image were used for statistics.

### Electrophysiological investigation of the structured neural networks

Using an identical microfluidic chip design interfaced with a custom-made microelectrode array (MEA), the electrophysiological activity of the structured neural networks and their response to KA stimulation were recorded through the MEA2100 in vitro Headstage, interface system and suit software (Multi Channel Systems; Reutlingen, Germany). Similar to the procedure described earlier, the MEA-interfaced microfluidic chips were coated with a combination of PLO and L15-laminin and seeded with 5×10^4^ NSCs per chamber (i.e. a total of 1.5×10^5^ per microfluidics chip), from which point synchronous differentiation and maturation of the NSCs into cortical neurons was carried out according to the manufacturer’s protocol until day 15. The electrophysiological activity of each MEA-interfaced structured neural network was recorded for a duration of 7 minutes immediately before KA stimulation, during stimulation, and at 24 hours post stimulation (10kHz sampling rate, 300Hz high-pass filter, with a ± 5 standard deviations upper- and lower spike detection threshold). For each of the two experimental groups (control neural networks vs LRRK2 neural networks) 6 MEA-interfaced neural networks were recorded and analyzed (3 controls, 3 KA stimulated). All raw data recordings are published in Mendeley Data Repository (doi: 10.17632/dnjv26msvk.4, doi: 10.17632/92568tpp39.4).

### Statistical analyses

Analyses were performed and visualized using Prism8 (GraphPad, California, United States). Quantifications from each assay were assessed for normality and homoscedasticity before being assigned a two-tailed parametric or non-parametric statistical test for comparing groups.

### Post-processing and analysis of electrophysiological data

Electrophysiological data analysis was performed with NeuroExplorer 4 (Nex Technologies, Colorado, United States) and MATLAB (MathWorks 2018, Massachusetts, United States). Following filtering and spike detection, the spikes were binned (1 sec) and the electrodes ordered according to chamber or channel of origin. Mean firing rates (MFRs) were estimated across conditions and recording time points as both total MFR and relative percentage deviation from baseline. Correlation was mapped using Pearson’s correlation coefficient r for concurrent spiking across 1-second spike binnings. Schemaballs (MATLAB central File Exchange, Komarov, Oleg, retrieved 2017.01.15) and heatmaps were used to display inter-electrode spike correlation ordered by chamber. Total network correlation was computed as mean r across electrodes per recording.

To illustrate the electrophysiological behavior of the LRRK2 neural networks in response to KA stimulation, a single electrode (electrode 63) was randomly selected for more detailed analysis. As before, spikes were detected by applying a simple thresholding approach to the electrophysiological data after bandpass filtering (passband: 300 Hz to 3 kHz). The spikes were then sorted using principal component analysis (PCA) to distinguish spikes from a single neuron that showed a period of enhanced firing rate accompanied by attenuation in spike amplitude, followed by an abrupt silencing. The firing rate was computed by convolving an alpha function kernel (α = 1 s^-1^, size = 10 s) with a spike train consisting of Dirac delta impulses at each spike time. The attenuation rates of the negative and trailing positive phases of the spikes were computed by fitting a line to the amplitude data during the period of high firing rate.

## 3. Results

Neural networks derived from both the control and LRRK2-mutated NSCs were successfully structured using the microfluidic chips (**Fig.1**, see also **Fig.6** is **STAR Method** for detailed microfluidic chip layout). Following 15 days of differentiation and maturation, immunocytochemistry confirmed the presence of neurons (MAP2), with neuron specific microtubules (beta-III tubulin) and mature axons (neurofilament heavy) containing both pre- and post-synaptic elements (Piccolo and PSD95, respectively), as well as expressing calmodulin-dependent protein kinase II (CamKII), a marker related to synaptic connectivity and long-term potentiation. Importantly, both kainic acid receptors (GRIK5) and AMPA receptors (GluR1) were also present (**Fig.1D, E and I**).

**Fig.1.**
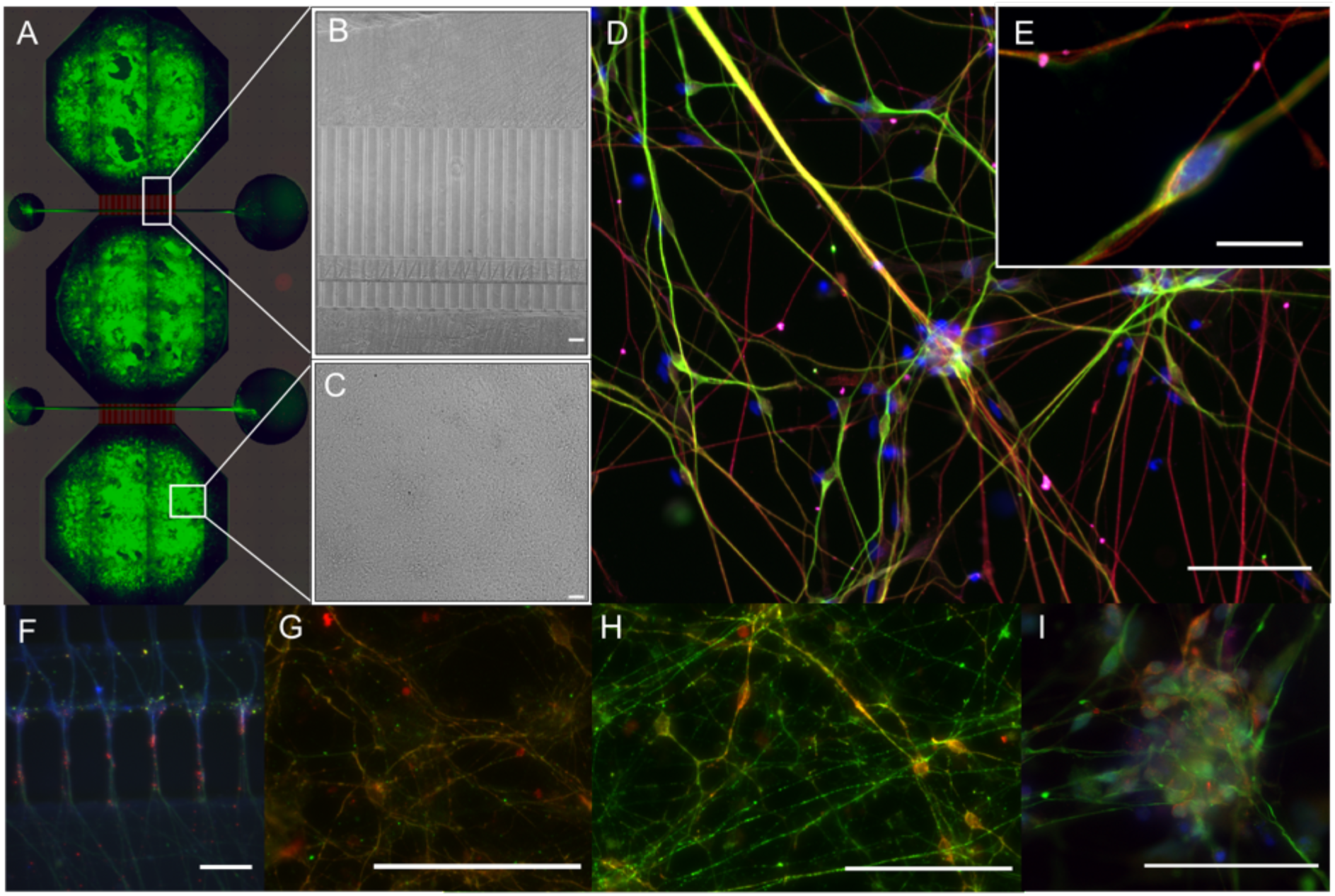
Structured neural networks. Following 15 days of NSC differentiation and maturation, immunocytochemistry confirmed the presence of mature neural networks in the microfluidic chips. **A**) Tiled image of a fluorescently labelled cortical neural network structured within a microfluidic chip, overlaid by a schematic of the design. **B**) Brightfield image of the developing neural network in a microfluidic chip, showing the area containing the axon tunnels, synaptic compartment and active zones, while **C**) Brightfield image from the cell chamber area. **D**) **F**luorescently labelled LRRK2 neural network with markers for neurons (MAP2, green), neuron specific microtubules (beta-III tubulin, red), and kainic acid receptors (GRIK5, magenta) together with the counterstain Hoechst (blue), with E) showing equivalent markers in a control neural network (10um scale bar). The remaining images show the neural networks fluorescently labelled with markers for **F**) neurons (MAP2, red) expressing (CaMKII, green), **G**) with presynaptic vesicles (Piccolo, green), postsynaptic densities (PSD95, red) and F-actin (Phalloidin, blue) expressed in the axon tunnels and synaptic area, **H**) neuronal specific microtubules (beta-III tubulin, red) together with CaMKII (green), and **I**) neurofilament heavy (green) together with AMPA receptors (red) and Hoechst counterstain. 50μm scale bars.

**Fig.6.**
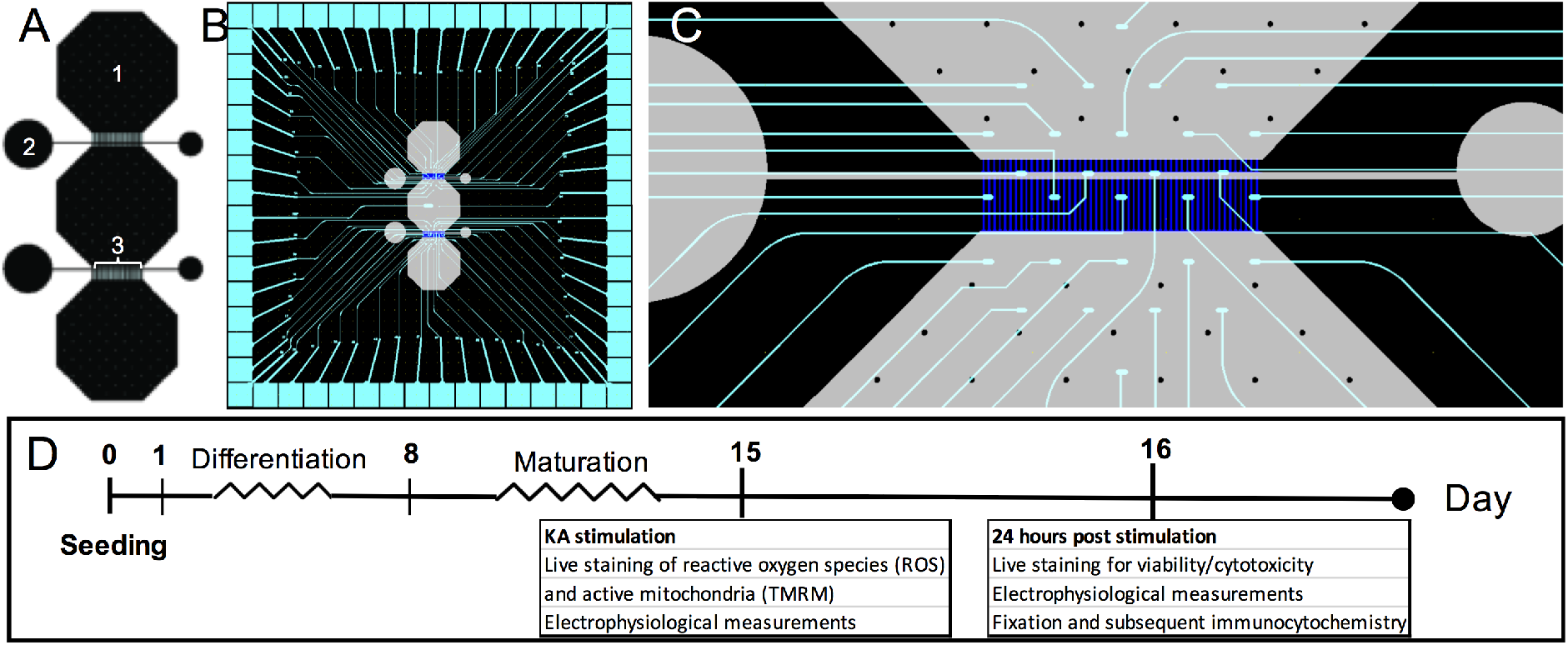
Experiment layout. **A**) Microfluidic chip design, with (1) indicating the top cell chamber/node, (2) the inlet/outlet connected via the synaptic compartment, and (3) the axonal/dendritic tunnels. Directionality of the inter-nodal connectivity is imposed on the network through different tunnel lengths, where mainly the axons from the top and bottom cell chambers connect to the dendrites and axons from the middle cell chamber in the two synaptic compartments. **B**) Outline of microfluidic chip interfaced with the custom made multielectrode array for electrophysiological investigation. **C**) Electrode layout in the area connecting the bottom and middle cell chamber through axonal/dendritic tunnels and the synaptic compartment. **D**) Timeline of the experiment, indicating the time points and assays used to assess the LRRK2 and control neural networks.

### Sublethal induction of overexcitation in structured neural networks by confined KA stimulation

Following a 30-minute KA stimulation targeting the neurons in the top cell chamber of a structured neural network, live microscopy of fluorescently labelled ROS production confirmed that the stimulation was successfully confined to the top cell chamber (**Fig.2A-E**). Furthermore, 24 hours post KA stimulation, the structured neural networks were fluorescently labelled with a live/dead viability assay kit, imaged and manually counted to determine whether the KA induced overexcitation was sublethal (**Fig.2F-H**). Two areas from each cell chamber were counted in two separate structured neural networks (a total of >2000 cells/network) for each condition (PBS condition vs KA) in each group (control vs LRRK2 neural networks), where the percentage of live cells found in each image was used for statistical analysis. To investigate whether there was a difference in cell viability between the KA-receiving and non-KA receiving chambers of the same neural networks, and between the different conditions in each group, a repeated measures two-way ANOVA was used. No statistically significant differences were found between chambers, indicating that the viability of the cells was not altered by the KA stimulation (p=0.3063). A statistically significant difference was found between the groups however (F_3,12_=25.3,7 p<0.0001). A post hoc Tukey’s multiple comparisons test found no significant difference between the KA stimulated and PBS condition within the same group (n_1_=n_2_=12 for both control and LRRK2 neural networks, with p=0.9996 and p=0.1921, respectively), confirming that the level of KA stimulation was sublethal. Interestingly, a significant difference was found for the PBS condition between the different neural populations (p=0.0004, DF=18.17), where a consistently higher percentage of viable cells was found in the LRRK2 neural networks compared to the control neural networks (**Fig.2H**).

**Fig.2.**
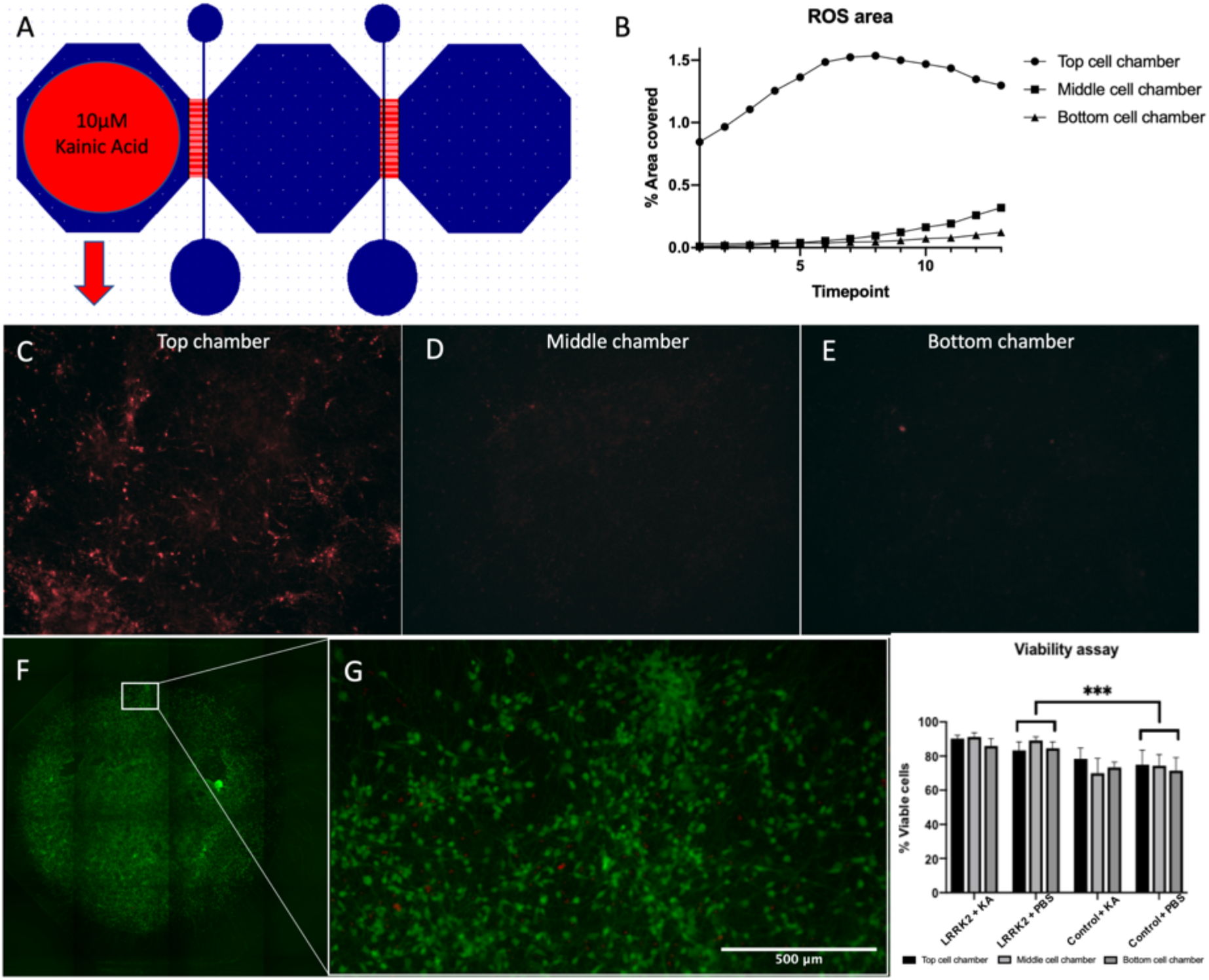
Confined sublethal kainic acid (KA) stimulation of neural networks in microfluidics chips. **A**) Microfluidic chip design, with a red circle indicating the top cell chamber used for confined KA stimulation (10μM). **B**) Line-graph where the reactive oxygen species (ROS) production of a structured neural network following a targeted KA stimulation has been analysed, with a clear difference between the ROS production in the top cell chamber compared to the two other chambers. Images (10X) were taken from each of the cell chambers every 5 minutes over the course of one hour following a KA stimulation. **C, D** and **E**) Representative images of fluorescently labelled ROS from each of the cell chambers, 45 minutes after the targeted KA stimulation. **F**) Tiled image of the middle cell chamber of a structured neural network fluorescently labelled with Calcein-Am (green) and Ethidium homodimer-1 (red) 24 hours post KA. **G**) Close-up of the area selectively chosen for analysis. **H**) Bar-graph showing the percentage of viable cells counted in each chamber, for each condition with standard deviation bars. A statistically significant difference was found between the control and the LRRK2 groups in the PBS condition (p=0.0004, N1=N2=12, DF=18.17) by post hoc Tukey’s multiple comparisons test, where N= the number of images counted. No significant difference was found between the KA and control condition within the same group, nor between the different chambers, demonstrating that the KA stimulation was sublethal as intended.

### Mitochondrial distribution in the axonal tunnels and synaptic compartments

Active mitochondria were successfully labelled with TMRM and could be visualized live in the structured neural networks using a Zeiss 510 META live confocal scanning microscope with a heated incubator. Single z-stacks containing all of the TMRM labelled mitochondria enclosed within a representative segment of the synaptic compartment were obtained at baseline for both the LRRK2 (n=6) and control (n=6) neural networks. Volumetric figures were produced for visualization of these data, and can be seen in **Fig.3A,B**. Height measures calculated from the z-stacks showed a statistically significant difference between the control and LRRK2 neural networks (two-tailed, independent samples t-test, t_10_=7.96, p<0.0001), with the LRRK2 neural networks containing mitochondria spanning on average over 3 times the height of the control neural networks (mean_KA_=35.38μm vs mean_healthy_=10.83μm) (**Fig.3A-E**). As the axonal tunnels restrict the movement of the mitochondria in the z-plane to a much greater extent than the synaptic compartment, they were chosen as the area for further imaging. The active mitochondria contained within a 145μm long segment of four axon tunnels (63X objective) were analyzed for the 6 control neural networks and the 5 LRRK2 neural networks, both at baseline and after KA or PBS addition. A statistically significant difference was found in the number of TMRM labelled mitochondria contained within the axon tunnels (Mann-Whitney U=659.5, N_1_=48, N_2_=40, p= 0.0114) between the control and LRRK2 neural networks at baseline, showing that the LRRK2 networks were characterized by the presence of a greater number of active mitochondria (median=164.5 vs 136) (**Fig.3F**). To investigate whether the number of active mitochondria was influenced by the KA stimulation, Wilcoxon matched-pairs signed ranks test was applied for the baseline and after KA stimulation timepoints in each group. A significant difference was found (pairs=24, p=0.0432) for the control neural networks, with a reduced number of active mitochondria measured after the KA stimulation. Although not statistically significant (pairs=24, p=0.1578), the same trend was observed for the KA stimulated condition in the LRRK2 neural networks, with fewer active mitochondria measured at the second timepoint than at baseline (**Fig.3I**). To control for potential degradation of the TMRM labelling over the time-course of the imaging, or a reduction of active mitochondria due to temperature changes moving from the incubator to the heated imaging-stage, or due to PBS-washing following the KA stimulation, networks (receiving only PBS) from each group were also imaged at the same timepoints. No significant difference was found between the two timepoints in the PBS condition for the control neural networks (pairs= 24, p=0.214) (**Fig.3H**). For the LRRK2 neural networks, a significant difference was found for the PBS condition, however, this time there were more active mitochondria at the latter timepoint (pairs=16, p=0.029). The mitochondrial sizes were also assessed for each condition; however, no significant differences were found between the baseline measures of mitochondrial size between the control and LRRK2 neural networks, nor between the baseline and KA stimulated condition in either group (data not shown).

**Fig.3.**
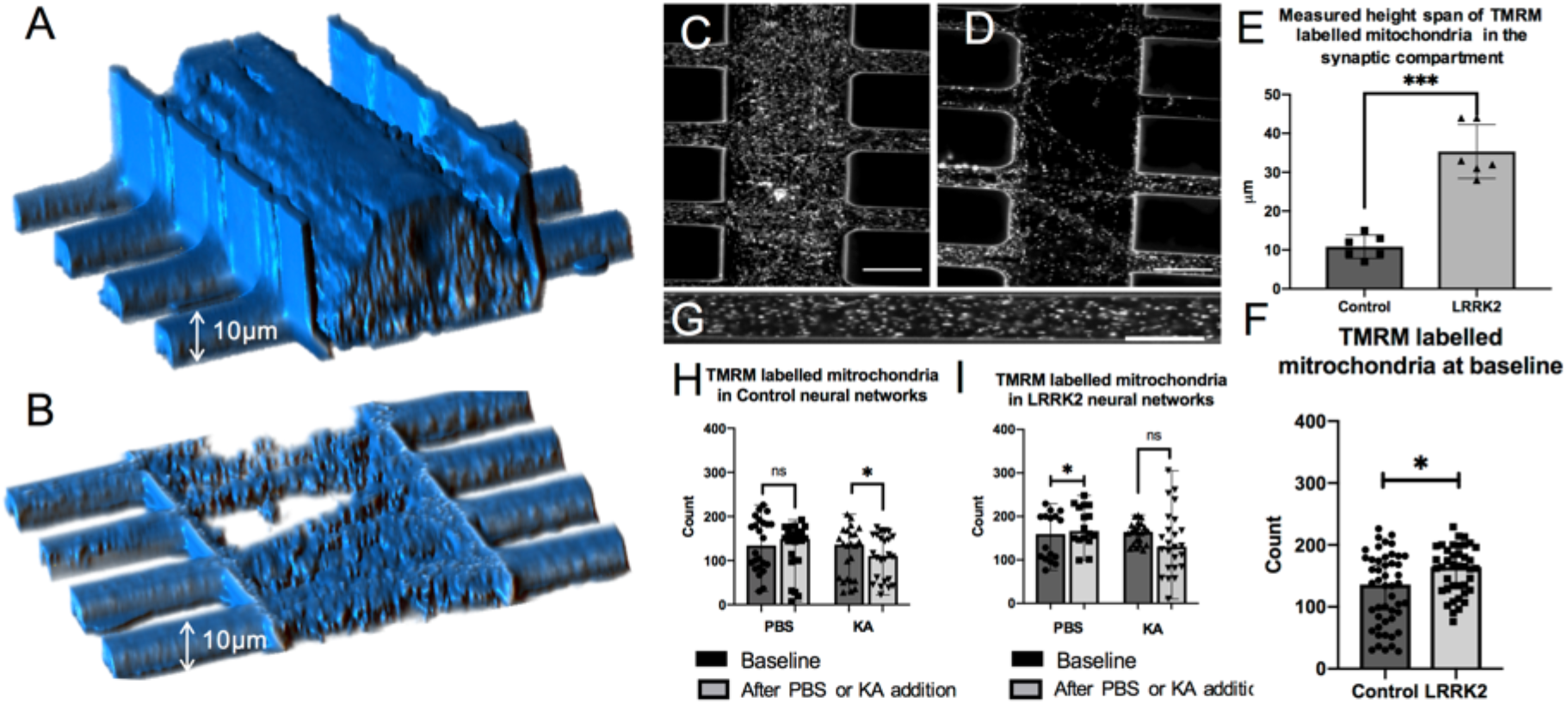
Mitochondrial distribution in the synaptic compartment and axon tunnels of the structured neural networks. The structured neural networks were labelled with TMRM, and the active mitochondria contained within 4 axonal tunnels were imaged live at two time points (baseline and after KA or PBS) for 3 neural networks from both groups, i.e. the control and LRRK2 neural networks. Additionally, image z-stacks were taken from the synaptic compartment at baseline for both groups to capture the height span of the active mitochondria. **A** and **B**) Volumetric view of the area containing fluorescently labelled mitochondria in the synaptic compartment from an **A**) LRRK2 neural network (height = 44μm) and a **B**) control neural network (height = 9μm). One of the z-slices from each of the stacks making up the volumetric figure in **A** and **B** are shown in **C** and **D**, respectively. Some autofluorescence in the PDMS walls of the microfluidic chips outline the structure of the synaptic compartment and axon tunnels (30μm scale bars). **E**) shows a bar-graph with scatter plots of the mean height in the synaptic compartment measured to contain fluorescently labelled mitochondria in each group, with standard deviation bars. An independent samples t-test showed that the LRRK2 neural networks contained active mitochondria within 3 times the height span of the control neural networks (p<0.0001, N=6) where N equals the number of networks investigated in each group. **G**) shows fluorescently labelled mitochondria contained within a single axonal tunnel in a control neural network, at baseline (20μm scale bar). **F**) Bar-graph with the median number of mitochondria counted within single axonal tunnels at baseline for both control and LRRK2 neural networks, with range bars and scatter plots. The LRRK2 neural networks displayed significantly more TMRM labelled mitochondria contained within the axonal tunnels compared to the control neural networks at baseline (p=0.0114, U=659.5, N_1_=48, N_2_=40) by Mann-Whitney U test. **H** and **I**) Bar-graphs with the median number of mitochondria measured at each timepoint, with range bars and scatter plots, for both the PBS and KA stimulated condition, within both groups. **H**) The control neural networks were found by Wilcoxon matched-pairs signed rank test to have significantly fewer active mitochondria after KA stimulation compared to the baseline (p=0.0432, pairs=24), while the LRRK2 neural networks **I**) showed the same trend without it being statistically significant. A statistically significant difference was found between the two timepoints in the PBS condition of the LRRK2 neural networks however (p=0.029, pairs=16), with more active mitochondria being measured after PBS addition.

Furthermore, to get an indication of whether the significant difference in number of TMRM labelled mitochondria observed at baseline between the control and LRRK2 neural networks (**Fig.3F**) was due to a greater mitochondrial content within each neurite in the LRRK2 neural networks compared to the control neural networks, a supplementary investigation of fluorescently labelled mitochondria was conducted in fixed samples from both groups. No statistically significant difference was found between the groups (p=0.4293, N_LRRK2_=8, N_control_=10) by independent samples t-test in this measure, indicating that the baseline difference in TMRM labelled mitochondria might be due to a larger number of neurites containing TMRM labelled mitochondria being visible, rather than a greater number of mitochondria being contained within each neurite, in the LRRK2 networks (**Supplementary Fig.S1**).

### Neurite morphology and internodal synaptic contacts

High-magnification microscopy images of the control neural networks fluorescently labelled with the anti-Piccolo antibody were used for morphological investigations of the neurites in the synaptic compartment, as well as for quantification of synaptic contacts through co-occurrence with the fluorescently labelled anti-PSD95 antibody, 24hours post KA stimulation (or PBS) (**Fig.4A-D**). Based on semi-automated image processing (see method section), a ratio of neuritic boutons/neurite was used to assess the morphology of the neurites, where a statistically significant difference was found between the PBS condition and the KA stimulated condition in the control neural networks (Mann-Whitney U=73, n_1_=22, n_2_=18, p=0.0004), with substantially fewer boutons observed after KA stimulation (median_PBS_=10.24 vs medianKA= 6.32). Furthermore, image analysis quantifying the number and size of co-occurring Piccolo and PSD95 labelling revealed a significant difference in size (μm) between the PBS and KA stimulated condition in the control neural networks (Mann-Whitney U=153.5, n_1_=26, n_2_=24, p=0.0017), with larger synaptic contact areas in the KA stimulated condition (median_PBS_=0.1915 vs median_KA_= 1.651), but no significant difference in number of contacts (median_PBS_=314.5 vs median_KA_=493.5, p=0.2504) (**Fig.4C, D**). Due to the high density of neurites contained within the synaptic compartment of the LRRK2 neural networks, z-stacks were taken to capture morphology of the neurites and the co-occurring Piccolo and PSD95 labelling (**Fig.S2 and S3**). The bottommost image slice from each stack was used for quantification of synaptic contacts through co-occurring Piccolo and PSD95 labelling (**Fig.4E,F**), while investigating the morphology of single neurites proved not feasible due to the overall compactness and density of the neurites in this group (**Fig.4G, Fig.S2**). No statistically significant differences were found in size (median_PBS_= 0.1190μm vs median_KA_= 0.1060μm) or number (median_PBS_=832 vs median_KA_=677) between the PBS and KA stimulated condition in the LRRK2 neural networks in co-occuring Piccolo and PSD95 (Mann-Whitney U test).

**Fig.4.**
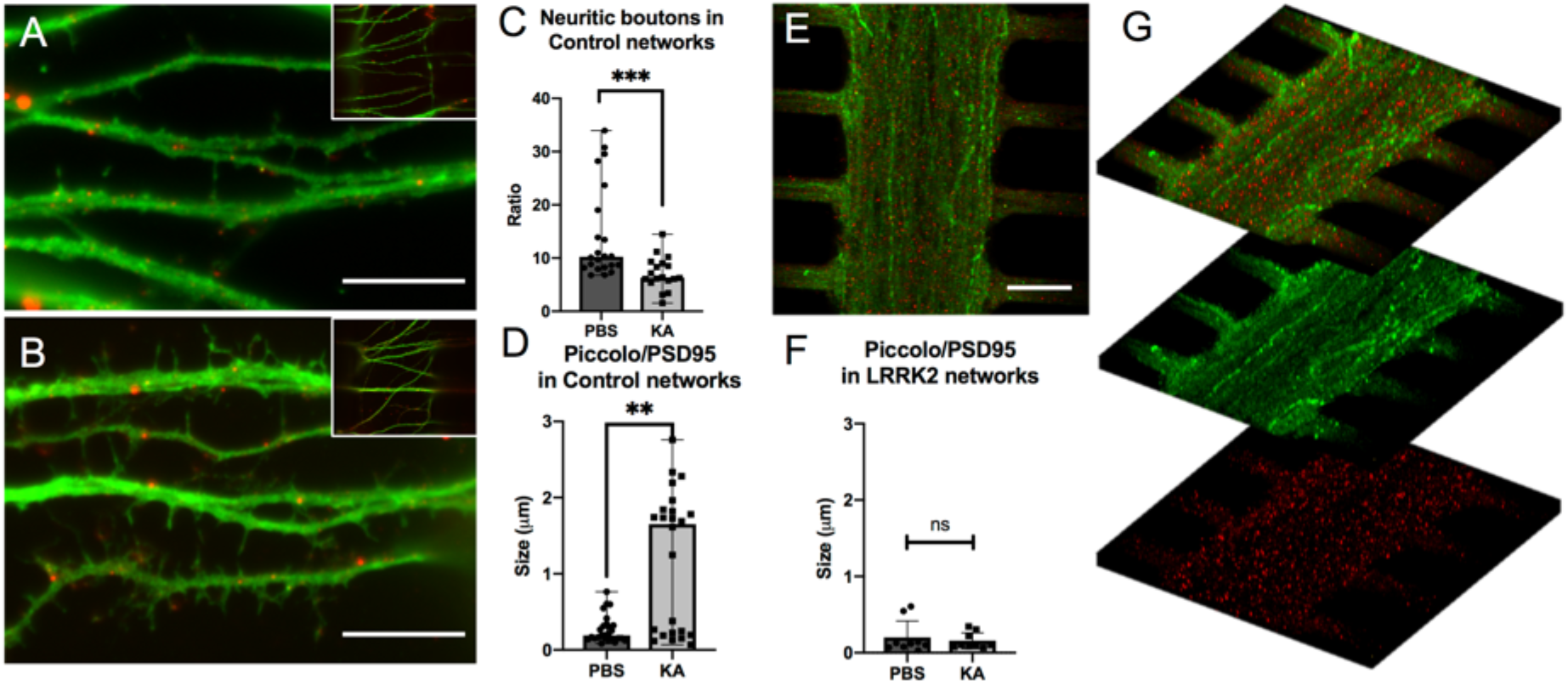
Neurite morphology and internodal synaptic contacts in the structured neural networks. Images were taken from the synaptic compartment area and show cortical neural networks fluorescently immunolabelled with Piccolo (green) and PSD95 (red). **A** and **B** Representative images from the two conditions (KA and PBS) in the control neural networks, where **A**) shows immunolabelling in the synaptic compartment of a KA stimulated neural network and **B**) from the synaptic compartment of a PBS neural network. Both images are enlarged (with representative full-view images in the top right corner) (10μm scale bars). **E**) Similarly, representative image from the KA stimulated condition in an LRRK2 neural network (20μm scale bar), illustrating the lower level of morphological detail available due to the density of neurites contained within the synaptic compartment. **C**) Bar-graph with the median ratio of neuritic boutons contained within the synaptic compartment for each condition in the control neural networks, with range bars. A statistically significant reduction of neuritic boutons was found in the KA stimulated condition (Mann-Whitney U=73, n_1_=22, n_2_=18, p=0.0004) compared to the PBS condition, where n equals the number of images analysed. **D** and **F**) Bar-graphs with the median size of the synaptic contacts (Piccolo/PSD95 co-occurrence) with range bars, measured within the synaptic compartment for both the control and LRRK2 neural networks, respectively. For the control neural networks, a statistically significant difference was found in the synaptic size measurements between the conditions (Mann-Whitney U=153.5, n_1_=26, n_2_=24, p=0.0017), with much larger areas of cooccurrence between Piccolo and PSD95 found at the neurites of the KA stimulated condition compared to the PBS condition. For the LRRK2 neural networks, no significant difference was found between the conditions in synaptic contact size. Furthermore, to illustrate the difference in neuritic density **G**) shows 10μm thick z-stack volume projections of the LRRK2 neural network in **E**, with Piccolo and PSD95 merged (top), followed by Piccolo (green) and PSD95 (red) alone.

### Electrophysiological activity of the structured neural networks

Electrophysiological measurements of the structured neural networks at baseline revealed a consistent trend in network activity and functional connectivity across all networks, both for control and LRRK2 neural networks, reflecting reproducible structure-function traits imposed on the networks through the physical structuring of the microfluidic chip. Higher mean firing rates (MFR) were consistently found at the top and bottom cell chambers relative to the middle cell chamber, and within-chamber (intra-nodal) correlations (Pearson’s correlation) were found to be consistently higher than between-chamber correlations, for both the LRRK2 and control neural networks (**Fig.S4**). Interestingly, the average MFR of the LRRK2 neural networks was far greater than that of the control neural networks at baseline (4.5 spikes/sec vs 2 spikes/sec) (**Fig.S4, Fig.5A**), while at the same time, the total network correlation was lower for the LRRK2 neural networks (r=0.085) than that of the control neural networks (r=0.12). Furthermore, average MFR and total network correlation measures from the PBS condition for both the LRRK2 and control neural networks (where some media was moved to create a flow barrier during the stimulation timepoint) follow each other quite closely (**Fig.5B, D**), with a comparable 33-39% drop in average MFR and a 43-44% increase in total network correlation at the 24 hours post-stimulation timepoint, relative to the baseline measures. The neural networks receiving KA produced differential responses however.

**Fig.5.**
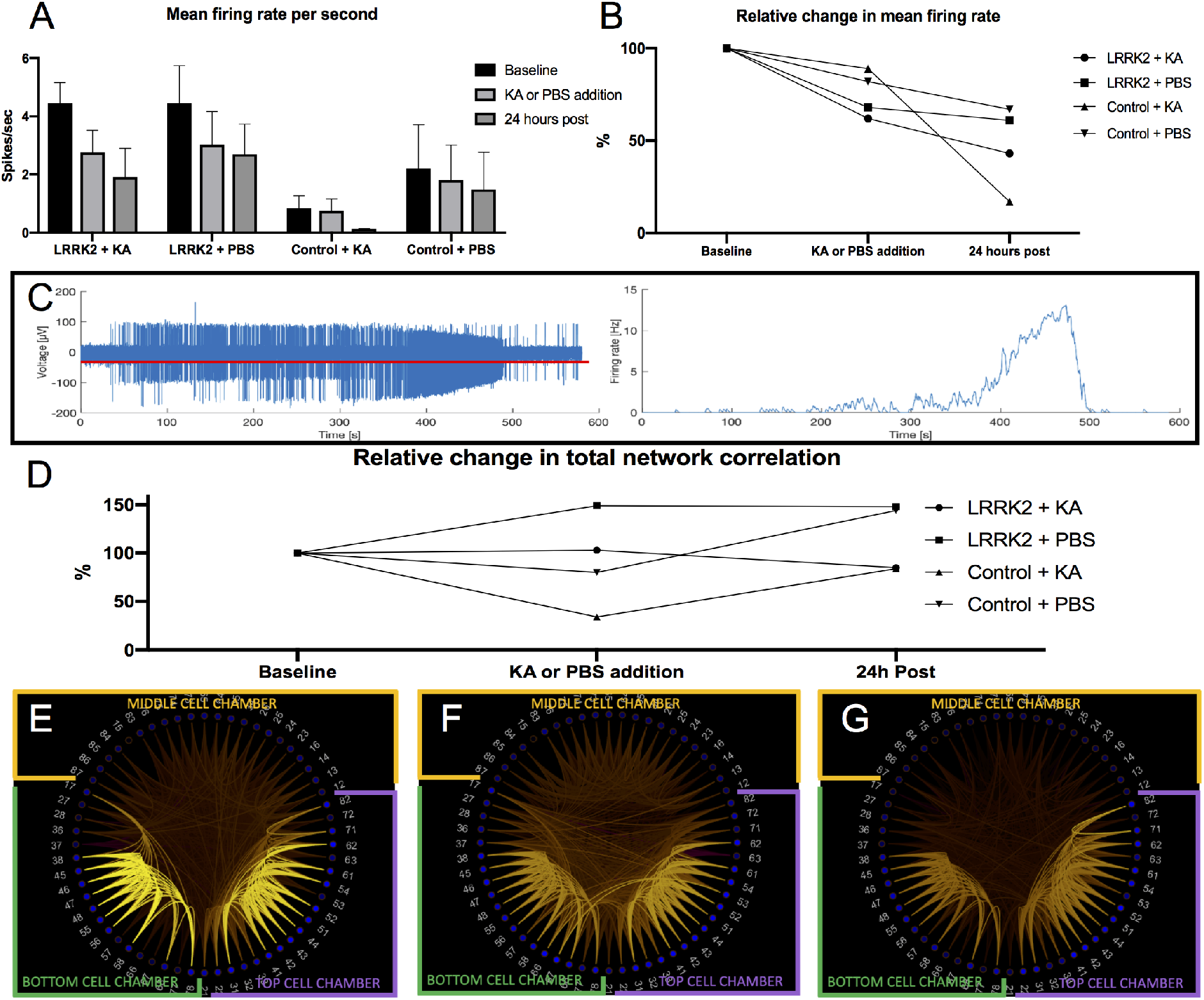
Electrophysiological measurements of the structured neural networks during baseline, KA stimulation, and 24 hours post stimulation. **A**) Bar graph of mean MFR with standard deviations measured for each group at the three different experimental timepoints (baseline, stimulation, 24hours post), with each group consisting of measurements from 3 different networks in each condition (PBS and KA) from both the control and LRRK2 group. The average MFR of the LRRK2 neural networks are consistently higher at all timepoints compared to the control neural networks, particularly at baseline. **B**) The same data as in **A**) but plotted as a line graph displaying the variation of the measures from the baseline condition. 24 hours post KA stimulation both the LRRK2 and control neural networks display a drop in MFR, however, the difference from the baseline is much larger for the control networks. **C**) Network overexcitation induced by the KA stimulation. The graph on the left shows the activity measured at a single electrode located in the top cell chamber of an LRRK2 neural network during the KA stimulation, with the red line indicating the threshold set for spike detection. The firing rate (Hz) profile of a single neuron recorded by the same electrode is plotted on the right (see Suppl.fig.4 for spike sorting details), where a drastic increase in firing can be observed from 350-500 seconds into the recording, followed by an abrupt activity drop. **D**) Line graph of the relative total network correlation change measured for each group at the three experimental timepoints in relation to the baseline values. At the 24 hours post KA or PBS addition timepoint, a similar increase in total network correlation change can be observed for the PBS condition of both the LRRK2 and control neural networks (+44-48%) relative to the baseline, while the KA stimulated networks display a similar decrease in total network correlation (−25-27%). **E-G**) Schemaball correlation maps of a representative LRRK2 neural network during the baseline, KA stimulation, and 24 hours post, respectively. Colour intensity of the lines interconnecting each channel indicates the correlation between their activity (from r 0-1), with 0 being black and 1 being bright yellow.

The KA addition successfully produced a transient overexcitation. The activity measured at a randomly chosen electrode (number 63) in the top chamber of an LRRK2 neural network during KA stimulation has been used to illustrate this in **Fig.5C** and **Fig.S4**., in which a transient period of substantially increased firing rate (i.e. from 2Hz to 14Hz) followed by an abrupt activity drop can be seen. Overall however, the average MFR during the entirety of the stimulation timepoint was reduced relative to the baseline, for both the LRRK2 and control neural networks (**Fig.5A**).

The relative MFR calculated from the baseline measures showed that the LRRK2 neural networks had a greater percentage drop in MFR during KA stimulation (38%) compared to the control neural networks (11%) (**Fig.5B**). At the same time, the LRRK2 neural networks exhibited a minor relative increase in total network correlation (3%) while the control neural networks displayed a relative reduction (64%) in total network correlation during the KA stimulation (**Fig.5D**). Although the overexcitation as demonstrated in **Fig.5C** was not observed in the PBS condition, the MFR and correlation measures from the networks in this condition also display large variations from the baseline during the stimulation timepoint, indicating that the simple procedure of creating a flow barrier alters the network activity.

24 hours later however, the neural networks in the PBS condition from both groups follow each other quite closely with comparable drops in MFRs and increases in total network correlations, as noted. The neural networks in the KA stimulated condition from both groups also display very similar, but reduced correlations at this timepoint, with the LRRK2 neural networks exhibiting a total network correlation of 85% and the control networks 83% relative to their baselines (**Fig.5D**). Interestingly, the KA stimulated networks differed widely in their relative MFR at this timepoint, where the LRRK2 neural networks exhibited a 57% drop, and the control neural networks exhibited a drastic 83% drop in MFR, relative to their baselines (**Fig.5B**). A representative KA stimulated LRRK2 neural network is used to visualize the overall network trend with correlation scheme balls from each of the timepoints in **Fig.5E-D**.

## 4. Discussion

### Baseline network activity measurements

One of the most striking findings in this report is the major difference in neuritic density observed at baseline between the neural networks carrying the G2019S LRRK2 mutation and the control neural networks (**Fig.3 and 5, Fig.S2, S3**). Several other studies have reported the involvement of the LRRK2 gene in neurite process morphology (West et al., 2005, Smith et al., 2005, MacLeod et al., 2006, Smith et al., 2006, Plowey et al., 2008, Dachsel et al., 2010, Gillardon, 2009, Meixner et al., 2011, Cherra et al., 2013, Habig et al., 2013, Sepulveda et al., 2013), where the specific G2019S PD-associated LRRK2 mutation has been found by most to increase kinase activity, resulting in a progressive reduction in neuritic length and branching (West et al., 2005, Smith et al., 2006, MacLeod et al., 2006, Plowey et al., 2008, Nguyen et al., 2011, Chan et al., 2011, Winner et al., 2011, Sanchez-Danes et al., 2012, Cherra et al., 2013, Reinhardt et al., 2013, Qing et al., 2017, Dagda et al., 2014, Greggio et al., 2006), with one exception demonstrating non-impaired neuritic morphology (Dachsel et al., 2010). Furthermore, knockdown and knock-out models resulting in LRRK2 deficiencies present with a progressive increase in neuritic length in some studies (MacLeod et al., 2006, Dachsel et al., 2010, Winner et al., 2011, Habig et al., 2013, Sepulveda et al., 2013), while others find the opposite (Gillardon, 2009, Meixner et al., 2011).

Interestingly, we provide the first evidence of increased neuritic density in human neurons with the G2019S mutation. This was demonstrated with immunocytochemistry of the neurites in the synaptic compartment (**Fig.4** and **Fig.S2**) and was further corroborated through several approaches using brightfield microscopy, Calcein-AM labelled cells (**Fig.S3**) and fluorescently labelled mitochondria (**Fig.3,S1**), all of which indicated a striking increase in neuritic density in LRRK2 neurons relative to the control neural networks. One of the studies mentioned above found the growth substrate to significantly influence the motility and outgrowth of neurites in culture (Sepulveda et al., 2013), with “fast-growth” coating substrates like laminin masking the effect of the mutation, and “slow-growth” substrates like poly-L-lysine enhancing them, something which might explain some of the variation between studies reported in the literature. Furthermore, structuring the neural networks with microfluidic chips in itself might enhance or reduce such traits (Dowell-Mesfin et al., 2004, Micholt et al., 2013). Thus, in our study, the combination of PLO and laminin as a growth substrate, together with structuring the neural networks in the microfluidic chip are likely to have influenced the resulting neuritic profile and may also partially explain some of the difference in relation to other studies. These factors do not explain the difference between our two groups, however, as the neural networks from both the LRRK2 and control group were treated in the exact same way, using identical protocols for coating, surface substrates, microfluidic chip structuring, cell differentiation and maturation, thus excluding differential culturing and handling as a potential source of variation between these populations. Furthermore, as the neural networks in the control and LRRK2 are derived from the same iPSC source, differences stemming from cell-line variability can also be excluded. Other possible influencing factors could be related to the source material, where most other studies use neural networks derived from rodents, and in some cases more region-specific cells related to the neurodegenerative pattern of PD, such as midbrain dopaminergic neurons, all of which might lead to variation in the results (West et al., 2005, Smith et al., 2006, MacLeod et al., 2006, Plowey et al., 2008, Nguyen et al., 2011, Chan et al., 2011, Winner et al., 2011, Sanchez-Danes et al., 2012, Cherra et al., 2013, Reinhardt et al., 2013, Qing et al., 2017, Dagda et al., 2014). Nevertheless, these apparent differences suggest that there are indeed subtleties in the mechanism underlying neuritic outgrowth and profile relating to the G2019S mutation that remain to be established.

As expected from the striking difference observed in neuritic profile between the networks, a significant difference in the average number of active mitochondria contained within the axonal tunnels in favor of the LRRK2 neural networks was also found (**Fig.3F**). Further investigation of fluorescently labelled mitochondria in fixed samples from each group revealed no significant difference in mitochondrial content within the cells of control versus LRRK2 neural networks (**Fig.S1**), suggesting that the observed difference in active TMRM labelled mitochondria is due to the larger volume of neurites being visible, rather than a larger number of active mitochondria within each neurite. Moreover, no significant difference in mitochondrial size was found at baseline between the two groups.

A greater volume of neurites should in theory provide greater opportunity for synapses to form and facilitate more efficient signal transduction, thus enabling greater structural connectivity. However, the LRRK2 neural networks displayed a lower total network correlation (r=0.085) at baseline compared to the control neural networks (r=0.12) (a difference which was reproduced in a second trial). Another fascinating observation is the major difference in baseline average MFR, where the LRRK2 neural networks consistently displayed about twice the MFR of the healthy neural networks (**Fig.5, Fig.S4**). In line with our observations, other recent studies have found the LRRK2-G2019S mutation to cause an increase in neural activity (Sweet et al., 2015, Volta et al., 2017, Matikainen-Ankney et al., 2016). Interestingly, one study has implicated increased neural activity as a pathogenic change preceding dendritic alteration in cortical neurons carrying this mutation (Plowey et al., 2014). Furthermore, G2019S-related hyperactivity has been shown (in vivo) to appear at an early point during development, which is proposed to affect the structure and function of the resulting striatal and other developing neural circuits, likely contributing to the progression of PD (Benson et al., 2018). Together this suggests a strong timescale-dependence of both the neural activity- and neuritic profile resulting from this mutation, where a difference in experimental timeframe might underlie some of the variation observed between studies of these features.

Taken together, our baseline measures show that the LRRK2 neural networks have an increase in neurites and in neural activity (MFR), which in turn is less well correlated across electrodes, relative to what is observed in the control neural networks. The elevated MFR and simultaneous low correlation represent *in vitro* neural network traits that are generally more prominent at very early time points, potentially pointing towards an overall impairment in LRRK2 neural network development. This notion of impaired development may be supported by another observation, i.e. that of differential growth cone profile at very early stages of LRRK2 neurons in culture compared to controls (**Fig.S6**). This may in turn explain what is observed as aberrant (and resultingly inefficient) network wiring in the microfluidic chips, where in the LRRK2 networks, large numbers of neurites can be seen crossing perpendicular to the axonal tunnels in the synaptic compartment (**Fig.4EG, Fig.S2, S3**). Furthermore, within the synaptic compartment, LRRK2 neurites show random outgrowth (**Fig.S3J**), compared to the directional, fasciculated outgrowth observed in the control networks (**Fig.S3E**). Taken together, these observations strongly suggest impaired axonal growth and guidance in the LRRK2 neurons. Another study also lends merit to this theory by finding altered neural growth cone morphology and number after knocking down LRRK2 (Habig et al., 2013). Furthermore, the altered neuritic profile might affect neurotransmission efficacy, resulting in less efficient signal propagation within the network, which may partially explain the observed increase in neural activity (assessed as MFR).

### Unveiling different network stress responses through transient overexcitation

The multiple-hit hypothesis of PD postulates that a combination of both genetic susceptibility and environmental factors may contribute to the onset and development of the disease (Carvey et al., 2006, Sulzer, 2007, Patrick et al., 2019). Based on this and the variability in disease progression in patients with LRRK2 associated PD, it is reasonable to assume that some phenotypic expression of the mutation may become apparent only following a significant or stressful challenge (Benson et al., 2018). Contrary to what one might expect, the neural networks carrying the G2019S LRRK2 mutation did not demonstrate a greater overall response to the transient overexcitation event compared to control neural networks. In fact, our one-off, sublethal, overexcitation event resulted in larger responses from the control neural networks in almost all measures, displaying a greater reduction in active mitochondria contained within the axonal tunnels immediately after the stimulation, as well as a more marked relative decrease in MFR, a more prominent alteration in neurite morphology, and greater synaptic remodeling 24 hours post stimulation, relative to the LRRK2 neural networks.

The highly significant difference found in neurite morphology in the control neural networks 24 hours post KA stimulation (**Fig.4A-C**) suggests neuritic remodeling in response to the transient excitatory stimulation event, with a retraction/reduction of boutons observed in the synaptic chambers in response to the KA overexcitation. At the same time, the number of synapses (co-occurrence of Piccolo and PSD95) in the control neural networks was not significantly altered, but the size of their overlapping area was, with much larger synaptic areas measured 24 hours after overexcitation (**Fig.4D**). This rapid, activity dependent alteration in spine morphology is likely an expression of a regular mechanism for converting short-term synaptic activity to long term lasting changes in connectivity and function. In line with our baseline measurements, the size range of the postsynaptic density is usually within 0.2-0.5μm, and can be localized to both spiny and non-spiny structures. Furthermore, this area contains both the kainate and AMPA receptors (Sheng, 2001), i.e. glutamatergic receptors targeted by our stimulation. During synaptic plasticity, the PSD increases in size in response to potentiation events, and the glutamate receptors contained within can be modulated by neural activity on a timescale from minutes to weeks (Sheng, 2001). In contrast to the observed neuritic alteration in the control neural networks, no significant alteration in synaptic number or size was found in the synaptic compartment of the LRRK2 mutated neural networks 24 hours after overexcitation, suggesting impaired synaptic plasticity. Some *in vivo* studies have implicated LRRK2 in the development, maturation and function of synapses, where the G2019S mutation has been shown to result in impaired long-term depression (Benson et al., 2018, Sweet et al., 2015). However, since our study investigates immediate and intermediate reactions (focused within the first 24 hours) to a single KA stimulation, no conclusions can be drawn about longer-term processes.

Furthermore, other studies have found increased vulnerability to oxidative stress, higher levels of mtDNA damage, and impaired mitochondrial movement as a result of the G2019S mutation, indicating compromised mitochondrial function (Cooper et al., 2012, Sanders et al., 2014, Hsieh et al., 2016, Schwab et al., 2017, Bose and Beal, 2019). Mitochondrial function alterations have in turn been shown to affect the plasticity of synapses and morphology of neurites as the availability of mitochondria is both essential and limiting for the support and maintenance of these structures (Li et al., 2004). In our study, both the control and the LRRK2 neural networks showed a reduction in the number of active mitochondria contained within the axon tunnels closest to the stimulated chamber immediately following the KA stimulation, however, only the change in the control networks was statistically significant. The energy status of the neuron greatly affects the motility and distribution of mitochondria (Saxton and Hollenbeck, 2012), and a retraction of the mitochondria towards the soma during or following an overexcitation event is to be expected as this corresponds to the location of greatest metabolic demand at the time. Loss of membrane potential in some of the mitochondria could also explain our results, as the TMRM labelling fades when this potential is disrupted. However, loss of mitochondrial membrane potential is the second observable step in classic excitotoxicity, preceded by ROS production, and followed by an opening of the permeability transition pore and induction of programmed cell death. Although our KA stimulation elicited a ROS response (as demonstrated in **Fig.2**), it did not result in any observable difference in cell viability 24 hours later, demonstrating a sublethal overexcitation as intended, effectively excluding classic excitotoxicity as the initiated process. A retraction of mitochondria towards the seat of greatest metabolic demand thus seems the most likely scenario. The statistically non-significant reduction in mitochondria observed in the LRRK2 neural networks following the overexcitation thus suggests an alteration in mitochondrial transport and/or function as a result of the G2019S mutation. The LRRK2 protein has been found to localize, among other sites, to mitochondrial structures where it interacts with a number of key regulators of mitochondrial fission/fusion, as well as to mitochondrial autophagy and motility (Biskup et al., 2006, Wang et al., 2012, Rosenbusch and Kortholt, 2016). Several other studies have found prominent signs of mitochondrial dysfunction relating both to size and distribution as a result of the G2019S mutation (MacLeod et al., 2006, Mortiboys et al., 2010, Li et al., 2014, Esteves et al., 2014, Yue et al., 2015, Singh et al., 2019), which strongly implies a role for LRRK2 in mitochondrial homeostasis. The initiating process leading to these mitochondrial alterations, however, are still elusive. Some possible factors are reduced mitochondrial biogenesis, impaired cytoskeletal elements and consequently cytoskeletal trafficking, direct functional and structural damage to the mitochondria by LRRK2 localized to the organelle, or impairment in the autophagic-lysosomal machinery resulting in accumulating damaged mitochondria (Yue et al., 2015, Tagliaferro and Burke, 2016, Franco-Iborra et al., 2018, Verma et al., 2018, Verma et al., 2017, Singh et al., 2019).

Intriguingly, 24 hours after the overexcitation, both the LRRK2 and the control neural networks displayed very well aligned decreases in relative total network correlations. This decrease becomes even more apparent when compared to the neural networks in the PBS condition from both groups, which drastically increased their correlations relative to the baseline at this point. This increase in PBS condition network correlations suggests that the neural networks were still maturing at the point of intervention, and that their natural trajectory was one towards greater correlation. The observed decrease in total network correlations is perhaps to be expected following such a stimulation as the overexcitation indiscriminately excites related connections that are both functional and non-functional, as well as produces both long-term potentiation and long-term depression of synapses based on coincidental activity at each connection. However, it is worth reiterating that these results were obtained after the neural networks were subjected to a single overexcitation event, while prolonged and/or repeated overexcitations would likely produce different results. The latter approach was beyond the scope of our current study. Nonetheless, both the LRRK2 and control neural networks also displayed a great drop in average MFR 24 hours after the overexcitation, where the relative drop was far greater for the control neural networks, demonstrating yet another impaired response in the LRRK2 neural networks comparted to the control neural networks.

In summary, in this study we provide the first evidence of increased neuritic density in structured, human neural networks carrying the G2019S LRRK2 mutation compared to control neural networks. At the same time, other network traits reported in the literature to result from this mutation are corroborated by our study, with the LRRK2 mutated neural networks displaying an increase in baseline neural activity (MFR) compared to the healthy control neural networks. Furthermore, eliciting a transient, confined, overexcitation through KA stimulation revealed different responses between the LRRK2 and control neural networks, both immediately after stimulation and 24 hours later. The control neural networks demonstrated a greater reduction in active mitochondria within the neurites immediately after the stimulation, as well a greater reduction in relative network activity (MFR), greater neuritic remodelling, and synaptic alterations, 24 hours later, compared to the LRRK2 neural networks.

It is thus clear that early signs of pathology relating to the G2019S LRRK2 mutation are captured within our chosen timeframe and reflected through both micro- and mesoscale investigations. Through a combination of neuroengineering, microfluidics, electrophysiology and fluorescence imaging strategies, we were able to selectively induce perturbations within interconnected network nodes and unveil these distinctions in a highly reproducible manner. Furthermore, as our neural networks were engineered from human cells with a homozygous Crispr-Cas9 inserted G2019S mutation, containing no artificially inserted gene or forced high or low levels of LRRK2, our *in vitro* setup builds on conditions genetically and physiologically more relevant to PD than networks engineered from model animal organisms. Moreover, as individuals homozygously carrying the G2019S mutation do not have more aggressive phenotypes compared to heterozygous carriers, there does not seem to be a dose-dependent increase in severity, and the results between studies of these two variants should be largely comparable (Ishihara et al., 2006, Lewis, 2019). Expanding the current platform to investigate these traits in even more detail and at greater length thus seems a logical next step. Several other mechanisms, which were out of the scope of this study, are of interest and might be involved in G2019S LRRK2 pathology, such as mitochondrial motility (Saxton and Hollenbeck, 2012, Singh et al., 2019), calcium overload and excitatory mitochondrial excitotoxicity (Verma et al., 2018, Verma et al., 2017), impaired synaptic endocytosis (Matta et al., 2012, Verstreken, 2017), alterations in the autophagosomal-lysosomal machinery (Sheehan and Yue, 2018, Verma et al., 2018, Pan et al., 2018, Singh et al., 2019), compromised AMPA receptor trafficking (Sweet et al., 2015), differential PKA regulation or altered Rab activity (Benson et al., 2018). Furthermore, as some of the investigated responses might also rely on cell-type specific mechanisms (Sweet et al., 2015, Pan et al., 2017, Benson et al., 2018), conducting a similar study with selectively vulnerable dopaminergic neurons or medium spiny neurons of the striatum might bring us even closer to understanding and unveiling pathological mechanisms of PD.

## Acknowledgements

This work was supported by The Department of Neuromedicine and Movement Science, Faculty of Medicine and Health Sciences, NTNU; The Liaison Committee for Education, Research and Innovation in Central Norway (Samarbeidsorganet HMN, NTNU); The Joint Research Committee between St Olav’s Hospital and the Faculty of Medicine and Health Sciences, NTNU; and the Research Council of Norway, Norwegian Micro- and Nano-Fabrication Facility, NorFab, project number 245963/F50, NTNU program for Enabling Technologies, (Nanotechnology). All imaging procedures requiring the use of the EVOS2 or the Zeiss 510 META live confocal scanning microscope were performed at the Cellular and Molecular Imaging Core Facility (CMIC), Norwegian University of Science and Technology (NTNU). CMIC is funded by the Faculty of Medicine at NTNU and Central Norwegian Regional Health Authority.

## Author contributions

V.D.V. Designed the study, conducted the experiments, collected and analyzed the data, and wrote the paper; R.W. Designed the study, the microfluidic chips and MEAs, contributed in execution of the experiments and data collection, contributed to data analysis and writing of the paper; O.H.R. Designed the study, performed the electrophysiology analysis, contributed to the data collection and writing of the methods; K.H. contributed with data analysis and writing of the methods; S.N provided supervision and valuable discussion in relation to the electrophysiology data analysis and techniques; A.S. Conceived and supervised the study, contributed to the study design, and critically reviewed the paper; I.S. Conceived and supervised the study, contributed to the study design, edited, and critically reviewed the paper.

## Data accessibility

All electrophysiology datasets have been deposited onto Mendeley and can be accessed through the following links:

doi: 10.17632/dnjv26msvk.4

doi: 10.17632/92568tpp39.4

## STAR Method

## Abbreviations

CamKII: calmodulin-dependent protein kinase II
iPSC: induced pluripotent stem cell
KA: kainic acid
LRRK2: leucine-rich repeat kinase 2
MEA: microelectrode array
MFR: mean firing rate
NGS: normal goat serum
NSC: neural stem cell
PBS: phosphate buffered saline
PCA: principal component analysis
PD: Parkinson’s disease
PDMS: polydimethylsiloxane
PFA: paraformaldehyde
PLO: poly-L-ornithine
PSD: post-synaptic density
ROS: reactive oxygen species
RT: room temperature
TMRM: tetramethylrhodiamine

## References

Baloyannis, S. J. 2006. Mitochondrial alterations in Alzheimer’s disease. J Alzheimers Dis, 9, 119–26.

Benson, D. L., Matikainen-Ankney, B. A., Hussein, A. & Huntley, G. W. 2018. Functional and behavioral consequences of Parkinson’s disease-associated LRRK2-G2019S mutation. Biochem Soc Trans, 46, 1697–1705.

Biskup, S., Moore, D. J., Celsi, F., Higashi, S., West, A. B., Andrabi, S. A., Kurkinen, K., Yu, S. W., Savitt, J. M., Waldvogel, H. J., Faull, R. L., Emson, P. C., Torp, R., Ottersen, O. P., Dawson, T. M. & Dawson, V. L. 2006. Localization of LRRK2 to membranous and vesicular structures in mammalian brain. Ann Neurol, 60, 557–69.

Bose, A. & Beal, M. F. 2016. Mitochondrial dysfunction in Parkinson’s disease. J Neurochem, 139 Suppl 1, 216–231.

Bose, A. & Beal, M. F. 2019. Mitochondrial dysfunction and oxidative stress in induced pluripotent stem cell models of Parkinson’s disease. Eur J Neurosci, 49, 525–532.

Bouhouche, A., Tibar, H., Ben El Haj, R., El Bayad, K., Razine, R., Tazrout, S., Skalli, A., Bouslam, N., Elouardi, L., Benomar, A., Yahyaoui, M. & Regragui, W. 2017. LRRK2 G2019S Mutation: Prevalence and Clinical Features in Moroccans with Parkinson’s Disease. Parkinsons Dis, 2017, 2412486.

Brizzee, K. R. 1987. Neurons numbers and dendritic extent in normal aging and Alzheimer’s disease. Neurobiol Aging, 8, 579–80.

Carvey, P. M., Punati, A. & Newman, M. B. 2006. Progressive dopamine neuron loss in Parkinson’s disease: the multiple hit hypothesis. Cell Transplant, 15, 239–50.

Caudle, W. M. & Zhang, J. 2009. Glutamate, excitotoxicity, and programmed cell death in Parkinson disease. Exp Neurol, 220, 230–3.

Chan, D., Citro, A., Cordy, J. M., Shen, G. C. & Wolozin, B. 2011. Rac1 protein rescues neurite retraction caused by G2019S leucine-rich repeat kinase 2 (LRRK2). J Biol Chem, 286, 16140–9.

Cheng, H. C., Ulane, C. M. & Burke, R. E. 2010. Clinical progression in Parkinson disease and the neurobiology of axons. Ann Neurol, 67, 715–25.

Cherra, S. J., 3RD, Steer, E., Gusdon, A. M., Kiselyov, K. & Chu, C. T. 2013. Mutant LRRK2 elicits calcium imbalance and depletion of dendritic mitochondria in neurons. Am J Pathol, 182, 474–84.

Cooper, O., Seo, H., Andrabi, S., Guardia-Laguarta, C., Graziotto, J., Sundberg, M., Mclean, J. R., Carrillo-Reid, L., Xie, Z., Osborn, T., Hargus, G., Deleidi, M., Lawson, T., Bogetofte, H., Perez-Torres, E., Clark, L., Moskowitz, C., Mazzulli, J., Chen, L., Volpicelli-Daley, L., Romero, N., Jiang, H., Uitti, R. J., Huang, Z., Opala, G., Scarffe, L. A., Dawson, V. L., Klein, C., Feng, J., Ross, O. A., Trojanowski, J. Q., Lee, V. M., Marder, K., Surmeier, D. J., Wszolek, Z. K., Przedborski, S., Krainc, D., Dawson, T. M. & Isacson, O. 2012. Pharmacological rescue of mitochondrial deficits in iPSC-derived neural cells from patients with familial Parkinson’s disease. Sci Transl Med, 4, 141ra90.

Dachsel, J. C., Behrouz, B., Yue, M., Beevers, J. E., Melrose, H. L. & Farrer, M. J. 2010. A comparative study of Lrrk2 function in primary neuronal cultures. Parkinsonism Relat Disord, 16, 650–5.

Dagda, R. K., Pien, I., Wang, R., Zhu, J., Wang, K. Z., Callio, J., Banerjee, T. D., Dagda, R. Y. & Chu, C. T. 2014. Beyond the mitochondrion: cytosolic PINK1 remodels dendrites through protein kinase A. J Neurochem, 128, 864–77.

Di Fonzo, A., Rohe, C. F., Ferreira, J., Chien, H. F., Vacca, L., Stocchi, F., Guedes, L., Fabrizio, E., Manfredi, M., Vanacore, N., Goldwurm, S., Breedveld, G., Sampaio, C., Meco, G., Barbosa, E., Oostra, B. A. & Bonifati, V. 2005. A frequent LRRK2 gene mutation associated with autosomal dominant Parkinson’s disease. Lancet, 365, 412–5.

Dowell-Mesfin, N. M., Abdul-Karim, M. A., Turner, A. M., Schanz, S., Craighead, H. G., Roysam, B., Turner, J. N. & Shain, W. 2004. Topographically modified surfaces affect orientation and growth of hippocampal neurons. J Neural Eng, 1, 78–90.

Esteves, A. R., Swerdlow, R. H. & Cardoso, S. M. 2014. LRRK2, a puzzling protein: insights into Parkinson’s disease pathogenesis. Exp Neurol, 261, 206–16.

Franco-Iborra, S., Vila, M. & Perier, C. 2018. Mitochondrial Quality Control in Neurodegenerative Diseases: Focus on Parkinson’s Disease and Huntington’s Disease. Front Neurosci, 12, 342.

Genc, B., Jara, J. H., Lagrimas, A. K., Pytel, P., Roos, R. P., Mesulam, M. M., Geula, C., Bigio, E. H. & Ozdinler, P. H. 2017. Apical dendrite degeneration, a novel cellular pathology for Betz cells in ALS. Sci Rep, 7, 41765.

Gilks, W. P., Abou-Sleiman, P. M., Gandhi, S., Jain, S., Singleton, A., Lees, A. J., Shaw, K., Bhatia, K. P., Bonifati, V., Quinn, N. P., Lynch, J., Healy, D. G., Holton, J. L., Revesz, T. & Wood, N. W. 2005. A common LRRK2 mutation in idiopathic Parkinson’s disease. Lancet, 365, 415–6.

Gillardon, F. 2009. Leucine-rich repeat kinase 2 phosphorylates brain tubulin-beta isoforms and modulates microtubule stability--a point of convergence in parkinsonian neurodegeneration? J Neurochem, 110, 1514–22.

Greggio, E., Jain, S., Kingsbury, A., Bandopadhyay, R., Lewis, P., Kaganovich, A., Van Der Brug, M. P., Beilina, A., Blackinton, J., Thomas, K. J., Ahmad, R., Miller, D. W., Kesavapany, S., Singleton, A., Lees, A., Harvey, R. J., Harvey, K. & Cookson, M. R. 2006. Kinase activity is required for the toxic effects of mutant LRRK2/dardarin. Neurobiol Dis, 23, 329–41.

Habig, K., Gellhaar, S., Heim, B., Djuric, V., Giesert, F., Wurst, W., Walter, C., Hentrich, T., Riess, O. & Bonin, M. 2013. LRRK2 guides the actin cytoskeleton at growth cones together with ARHGEF7 and Tropomyosin 4. Biochim Biophys Acta, 1832, 2352–67.

Habig, K., Walter, M., Poths, S., Riess, O. & Bonin, M. 2008. RNA interference of LRRK2-microarray expression analysis of a Parkinson’s disease key player. Neurogenetics, 9, 83–94.

Healy, D. G., Falchi, M., O’sullivan, S. S., Bonifati, V., Durr, A., Bressman, S., Brice, A., Aasly, J., Zabetian, C. P., Goldwurm, S., Ferreira, J. J., Tolosa, E., Kay, D. M., Klein, C., Williams, D. R., Marras, C., Lang, A. E., Wszolek, Z. K., Berciano, J., Schapira, A. H., Lynch, T., Bhatia, K. P., Gasser, T., Lees, A. J. & Wood, N. W. 2008. Phenotype, genotype, and worldwide genetic penetrance of LRRK2-associated Parkinson’s disease: a case-control study. Lancet Neurol, 7, 583–90.

Hsieh, C. H., Shaltouki, A., Gonzalez, A. E., Bettencourt Da Cruz, A., Burbulla, L. F., St Lawrence, E., Schule, B., Krainc, D., Palmer, T. D. & Wang, X. 2016. Functional Impairment in Miro Degradation and Mitophagy Is a Shared Feature in Familial and Sporadic Parkinson’s Disease. Cell Stem Cell, 19, 709–724.

Ishihara, L., Warren, L., Gibson, R., Amouri, R., Lesage, S., Durr, A., Tazir, M., Wszolek, Z. K., Uitti, R. J., Nichols, W. C., Griffith, A., Hattori, N., Leppert, D., Watts, R., Zabetian, C. P., Foroud, T. M., Farrer, M. J., Brice, A., Middleton, L. & Hentati, F. 2006. Clinical features of Parkinson disease patients with homozygous leucine-rich repeat kinase 2 G2019S mutations. Arch Neurol, 63, 1250–4.

Jaleel, M., Nichols, R. J., Deak, M., Campbell, D. G., Gillardon, F., Knebel, A. & Alessi, D. R. 2007. LRRK2 phosphorylates moesin at threonine-558: characterization of how Parkinson’s disease mutants affect kinase activity. Biochem J, 405, 307–17.

Kachergus, J., Mata, I. F., Hulihan, M., Taylor, J. P., Lincoln, S., Aasly, J., Gibson, J. M., Ross, O. A., Lynch, T., Wiley, J., Payami, H., Nutt, J., Maraganore, D. M., Czyzewski, K., Styczynska, M., Wszolek, Z. K., Farrer, M. J. & Toft, M. 2005. Identification of a novel LRRK2 mutation linked to autosomal dominant parkinsonism: evidence of a common founder across European populations. Am J Hum Genet, 76, 672–80.

Lesage, S., Ibanez, P., Lohmann, E., Pollak, P., Tison, F., Tazir, M., Leutenegger, A. L., Guimaraes, J., Bonnet, A. M., Agid, Y., Durr, A. & Brice, A. 2005. G2019S LRRK2 mutation in French and North African families with Parkinson’s disease. Ann Neurol, 58, 784–7.

Lewis, P. A. 2019. Leucine rich repeat kinase 2: a paradigm for pleiotropy. J Physiol, 597, 3511–3521.

Li, J. Q., Tan, L. & Yu, J. T. 2014. The role of the LRRK2 gene in Parkinsonism. Mol Neurodegener, 9, 47.

Li, Z., Okamoto, K., Hayashi, Y. & Sheng, M. 2004. The importance of dendritic mitochondria in the morphogenesis and plasticity of spines and synapses. Cell, 119, 873–87.

Macleod, D., Dowman, J., Hammond, R., Leete, T., Inoue, K. & Abeliovich, A. 2006. The familial Parkinsonism gene LRRK2 regulates neurite process morphology. Neuron, 52, 587–93.

Matikainen-Ankney, B. A., Kezunovic, N., Mesias, R. E., Tian, Y., Williams, F. M., Huntley, G. W. & Benson, D. L. 2016. Altered Development of Synapse Structure and Function in Striatum Caused by Parkinson’s Disease-Linked LRRK2-G2019S Mutation. J Neurosci, 36, 7128–41.

Matta, S., Van Kolen, K., Da Cunha, R., Van Den Bogaart, G., Mandemakers, W., Miskiewicz, K., De Bock, P. J., Morais, V. A., Vilain, S., Haddad, D., Delbroek, L., Swerts, J., Chavez-Gutierrez, L., Esposito, G., Daneels, G., Karran, E., Holt, M., Gevaert, K., Moechars, D. W., De Strooper, B. & Verstreken, P. 2012. LRRK2 controls an EndoA phosphorylation cycle in synaptic endocytosis. Neuron, 75, 1008–21.

Meixner, A., Boldt, K., Van Troys, M., Askenazi, M., Gloeckner, C. J., Bauer, M., Marto, J. A., Ampe, C., Kinkl, N. & Ueffing, M. 2011. A QUICK screen for Lrrk2 interaction partners--leucine-rich repeat kinase 2 is involved in actin cytoskeleton dynamics. Mol Cell Proteomics, 10, M110.001172.

Micholt, L., Gartner, A., Prodanov, D., Braeken, D., Dotti, C. G. & Bartic, C. 2013. Substrate topography determines neuronal polarization and growth in vitro. PLoS One, 8, e66170.

Moore, D. J. 2008. The biology and pathobiology of LRRK2: implications for Parkinson’s disease. Parkinsonism Relat Disord, 14 Suppl 2, S92–8.

Mortiboys, H., Johansen, K. K., Aasly, J. O. & Bandmann, O. 2010. Mitochondrial impairment in patients with Parkinson disease with the G2019S mutation in LRRK2. Neurology, 75, 2017–20.

Nguyen, H. N., Byers, B., Cord, B., Shcheglovitov, A., Byrne, J., Gujar, P., Kee, K., Schule, B., Dolmetsch, R. E., Langston, W., Palmer, T. D. & Pera, R. R. 2011. LRRK2 mutant iPSC-derived DA neurons demonstrate increased susceptibility to oxidative stress. Cell Stem Cell, 8, 267–80.

Nichols, W. C., Pankratz, N., Hernandez, D., Paisan-Ruiz, C., Jain, S., Halter, C. A., Michaels, V. E., Reed, T., Rudolph, A., Shults, C. W., Singleton, A. & Foroud, T. 2005. Genetic screening for a single common LRRK2 mutation in familial Parkinson’s disease. Lancet, 365, 410–2.

Ozelius, L. J., Senthil, G., Saunders-Pullman, R., Ohmann, E., Deligtisch, A., Tagliati, M., Hunt, A. L., Klein, C., Henick, B., Hailpern, S. M., Lipton, R. B., Soto-Valencia, J., Risch, N. & Bressman, S. B. 2006. LRRK2 G2019S as a cause of Parkinson’s disease in Ashkenazi Jews. N Engl J Med, 354, 424–5.

Pan, P. Y., Li, X., Wang, J., Powell, J., Wang, Q., Zhang, Y., Chen, Z., Wicinski, B., Hof, P., Ryan, T. A. & Yue, Z. 2017. Parkinson’s Disease-Associated LRRK2 Hyperactive Kinase Mutant Disrupts Synaptic Vesicle Trafficking in Ventral Midbrain Neurons. J Neurosci, 37, 11366–11376.

Pan, P. Y., Zhu, Y., Shen, Y. & Yue, Z. 2018. Crosstalk between presynaptic trafficking and autophagy in Parkinson’s disease. Neurobiol Dis.

Patrick, K. L., Bell, S. L., Weindel, C. G. & Watson, R. O. 2019. Exploring the “Multiple-Hit Hypothesis” of Neurodegenerative Disease: Bacterial Infection Comes Up to Bat. Front Cell Infect Microbiol, 9, 138.

Plowey, E. D., Cherra, S. J., 3RD, Liu, Y. J. & Chu, C. T. 2008. Role of autophagy in G2019S-LRRK2-associated neurite shortening in differentiated SH-SY5Y cells. J Neurochem, 105, 1048–56.

Plowey, E. D., Johnson, J. W., Steer, E., Zhu, W., Eisenberg, D. A., Valentino, N. M., Liu, Y. J. & Chu, C. T. 2014. Mutant LRRK2 enhances glutamatergic synapse activity and evokes excitotoxic dendrite degeneration. Biochim Biophys Acta, 1842, 1596–603

Qing, X., Walter, J., Jarazo, J., Arias-Fuenzalida, J., Hillje, A. L. & Schwamborn, J. C. 2017. CRISPR/Cas9 and piggyBac-mediated footprint-free LRRK2-G2019S knock-in reveals neuronal complexity phenotypes and alpha-Synuclein modulation in dopaminergic neurons. Stem Cell Res, 24, 44–50.

Reinhardt, P., Schmid, B., Burbulla, L. F., Schondorf, D. C., Wagner, L., Glatza, M., Hoing, S., Hargus, G., Heck, S. A., Dhingra, A., Wu, G., Muller, S., BRockmann, K., Kluba, T., Maisel, M., Kruger, R., Berg, D., Tsytsyura, Y., Thiel, C. S., Psathaki, O. E., Klingauf, J., Kuhlmann, T., Klewin, M., Muller, H., Gasser, T., Scholer, H. R. & Sterneckert, J. 2013. Genetic correction of a LRRK2 mutation in human iPSCs links parkinsonian neurodegeneration to ERK-dependent changes in gene expression. Cell Stem Cell, 12, 354–67.

Rosenbusch, K. E. & Kortholt, A. 2016. Activation Mechanism of LRRK2 and Its Cellular Functions in Parkinson’s Disease. Parkinsons Dis, 2016, 7351985.

Sanchez-Danes, A., Richaud-Patin, Y., Carballo-Carbajal, I., Jimenez-Delgado, S., Caig, C., Mora, S., Di Guglielmo, C., Ezquerra, M., Patel, B., Giralt, A., Canals, J. M., Memo, M., Alberch, J., Lopez-Barneo, J., Vila, M., Cuervo, A. M., Tolosa, E., Consiglio, A. & Raya, A. 2012. Disease-specific phenotypes in dopamine neurons from human iPS-based models of genetic and sporadic Parkinson’s disease. EMBO Mol Med, 4, 380–95.

Sanders, L. H., Laganiere, J., Cooper, O., Mak, S. K., Vu, B. J., Huang, Y. A., Paschon, D. E., Vangipuram, M., Sundararajan, R., Urnov, F. D., Langston, J. W., Gregory, P. D., Zhang, H. S., Greenamyre, J. T., Isacson, O. & Schule, B. 2014. LRRK2 mutations cause mitochondrial DNA damage in iPSC-derived neural cells from Parkinson’s disease patients: reversal by gene correction. Neurobiol Dis, 62, 381–6.

Sasaki, S. & Iwata, M. 2007. Mitochondrial alterations in the spinal cord of patients with sporadic amyotrophic lateral sclerosis. J Neuropathol Exp Neurol, 66, 10–6.

Saxton, W. M. & Hollenbeck, P. J. 2012. The axonal transport of mitochondria. J Cell Sci, 125, 2095–104.

Schwab, A. J., Sison, S. L., Meade, M. R., Broniowska, K. A., Corbett, J. A. & Ebert, A. D. 2017. Decreased Sirtuin Deacetylase Activity in LRRK2 G2019S iPSC-Derived Dopaminergic Neurons. Stem Cell Reports, 9, 1839–1852.

Sepulveda, B., Mesias, R., Li, X., Yue, Z. & Benson, D. L. 2013. Short-and long-term effects of LRRK2 on axon and dendrite growth. PLoS One, 8, e61986.

Sheehan, P. & Yue, Z. 2018. Deregulation of autophagy and vesicle trafficking in Parkinson’s disease. Neurosci Lett.

Sheng, M. H.-T. 2001. The postsynaptic specialization In: Cowan, W. M., SÜdhof, T. C. & Stevens, C. F. (eds.) Synapses. The John Hopkins University Press.

Singh, A., Zhi, L. & Zhang, H. 2019. LRRK2 and mitochondria: Recent advances and current views. Brain Res, 1702, 96–104.

Smith, W. W., Pei, Z., Jiang, H., Dawson, V. L., Dawson, T. M. & Ross, C. A. 2006. Kinase activity of mutant LRRK2 mediates neuronal toxicity. Nat Neurosci, 9, 1231–3.

Smith, W. W., Pei, Z., Jiang, H., Moore, D. J., Liang, Y., West, A. B., Dawson, V. L., Dawson, T. M. & Ross, C. A. 2005. Leucine-rich repeat kinase 2 (LRRK2) interacts with parkin, and mutant LRRK2 induces neuronal degeneration. Proc Natl Acad Sci U S A, 102, 18676–81.

Steger, M., Tonelli, F., Ito, G., Davies, P., Trost, M., Vetter, M., Wachter, S., Lorentzen, E., Duddy, G., Wilson, S., Baptista, M. A., Fiske, B. K., Fell, M. J., Morrow, J. A., Reith, A. D., Alessi, D. R. & Mann, M. 2016. Phosphoproteomics reveals that Parkinson’s disease kinase LRRK2 regulates a subset of Rab GTPases. Elife, 5.

Stephens, B., Mueller, A. J., Shering, A. F., Hood, S. H., Taggart, P., Arbuthnott, G. W., Bell, J. E., Kilford, L., Kingsbury, A. E., Daniel, S. E. & Ingham, C. A. 2005. Evidence of a breakdown of corticostriatal connections in Parkinson’s disease. Neuroscience, 132, 741–54.

Sulzer, D. 2007. Multiple hit hypotheses for dopamine neuron loss in Parkinson’s disease. Trends Neurosci, 30, 244–50.

Sweet, E. S., Saunier-Rebori, B., Yue, Z. & Blitzer, R. D. 2015. The Parkinson’s Disease-Associated Mutation LRRK2-G2019S Impairs Synaptic Plasticity in Mouse Hippocampus. J Neurosci, 35, 11190–5.

Tagliaferro, P. & Burke, R. E. 2016. Retrograde Axonal Degeneration in Parkinson Disease. J Parkinsons Dis, 6, 1–15.

Trinh, J., Farrer, M., Ross, A. O. & Guella, I. 2006. LRRK2-Related Parkinsons Disease. In: Adam MP, Ardinger HH & Ra, P. (eds.). Seattle (WA): University of Washington, Seattle: GeneReviews® [Internet].

Van De Wijdeven, R., Ramstad, O. H., Bauer, U. S., Halaas, O., Sandvig, A. & Sandvig, I. 2018. Structuring a multi-nodal neural network in vitro within a novel design microfluidic chip. Biomed Microdevices, 20, 9.

Van De Wijdeven, R., Ramstad, O. H., Valderhaug, V. D., Kollensperger, P., Sandvig, A., Sandvig, I. & Halaas, O. 2019. A novel lab-on-chip platform enabling axotomy and neuromodulation in a multi-nodal network. Biosens Bioelectron, 140, 111329.

Verma, M., Callio, J., Otero, P. A., Sekler, I., Wills, Z. P. & Chu, C. T. 2017. Mitochondrial Calcium Dysregulation Contributes to Dendrite Degeneration Mediated by PD/LBD-Associated LRRK2 Mutants. J Neurosci, 37, 11151–11165.

Verma, M., Wills, Z. & Chu, C. T. 2018. Excitatory Dendritic Mitochondrial Calcium Toxicity: Implications for Parkinson’s and Other Neurodegenerative Diseases. Front Neurosci, 12, 523.

Verstreken, P. (ed.) 2017. Parkinson’s Disease: Molecular Mechanisms Underlying Pathology Academic Press.

Volta, M., Beccano-Kelly, D. A., Paschall, S. A., Cataldi, S., Macisaac, S. E., Kuhlmann, N., Kadgien, C. A., Tatarnikov, I., Fox, J., Khinda, J., Mitchell, E., Bergeron, S., Melrose, H., Farrer, M. J. & Milnerwood, A. J. 2017. Initial elevations in glutamate and dopamine neurotransmission decline with age, as does exploratory behavior, in LRRK2 G2019S knock-in mice. Elife, 6.

Wang, X., Yan, M. H., Fujioka, H., Liu, J., Wilson-Delfosse, A., Chen, S. G., Perry, G., Casadesus, G. & Zhu, X. 2012. LRRK2 regulates mitochondrial dynamics and function through direct interaction with DLP1. Hum Mol Genet, 21, 1931–44.

West, A. B., Moore, D. J., Biskup, S., Bugayenko, A., Smith, W. W., Ross, C. A., Dawson, V. L. & Dawson, T. M. 2005. Parkinson’s disease-associated mutations in leucine-rich repeat kinase 2 augment kinase activity. Proc Natl Acad Sci U S A, 102, 16842–7.

Winner, B., Melrose, H. L., Zhao, C., Hinkle, K. M., Yue, M., Kent, C., Braithwaite, A. T., Ogholikhan, S., Aigner, R., Winkler, J., Farrer, M. J. & Gage, F. H. 2011. Adult neurogenesis and neurite outgrowth are impaired in LRRK2 G2019S mice. Neurobiol Dis, 41, 706–16.

Yue, M., Hinkle, K. M., Davies, P., Trushina, E., Fiesel, F. C., Christenson, T. A., Schroeder, A. S., Zhang, L., Bowles, E., Behrouz, B., Lincoln, S. J., Beevers, J. E., Milnerwood, A. J., Kurti, A., Mclean, P. J., Fryer, J. D., Springer, W., Dickson, D. W., Farrer, M. J. & Melrose, H. L. 2015. Progressive dopaminergic alterations and mitochondrial abnormalities in LRRK2 G2019S knock-in mice. Neurobiol Dis, 78, 172–95.

Zhao, J., Hermanson, S. B., Carlson, C. B., Riddle, S. M., Vogel, K. W., Bi, K. & Nichols, R. J. 2012. Pharmacological inhibition of LRRK2 cellular phosphorylation sites provides insight into LRRK2 biology. Biochem Soc Trans, 40, 1158–62.

