## Supplementary document for "Structural and functional alterations associated with the LRRK2 G2019S mutation revealed in structured human neural networks"

### Supplementary figures

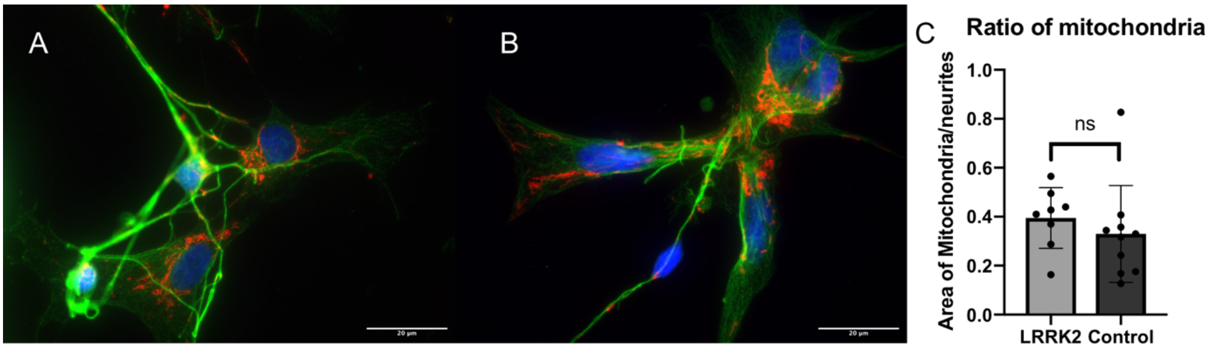


**Fig.S1. Supplementary investigation of mitochondria A**) shows a LRRK2 neural network labelled with fluorescent mitochondria (red (ab3298)), total-alpha synuclein (green) and hoechst conterstain (blue). **B**) shows an equivalent image of a fluorescently labelled control neural network (100X objective). **C**) shows a bar graph of the ratio calculation of mitochondria contained within samples from both the LRRK2 and Control neural networks at baseline, with example images used for the calculation displayed in panel **A**) and **B**). No statistically significant difference was found between the groups in mitochondrial content (ratio = area of fluorescently labelled mitochondria/ area of fluorescently labelled alpha-synuclein) (p=0.4293) by an unpaired t-test. Each dot represents a different sample (100X image) investigated, where $N_{LRRK2}$=8, and $N_{Control}$=10.

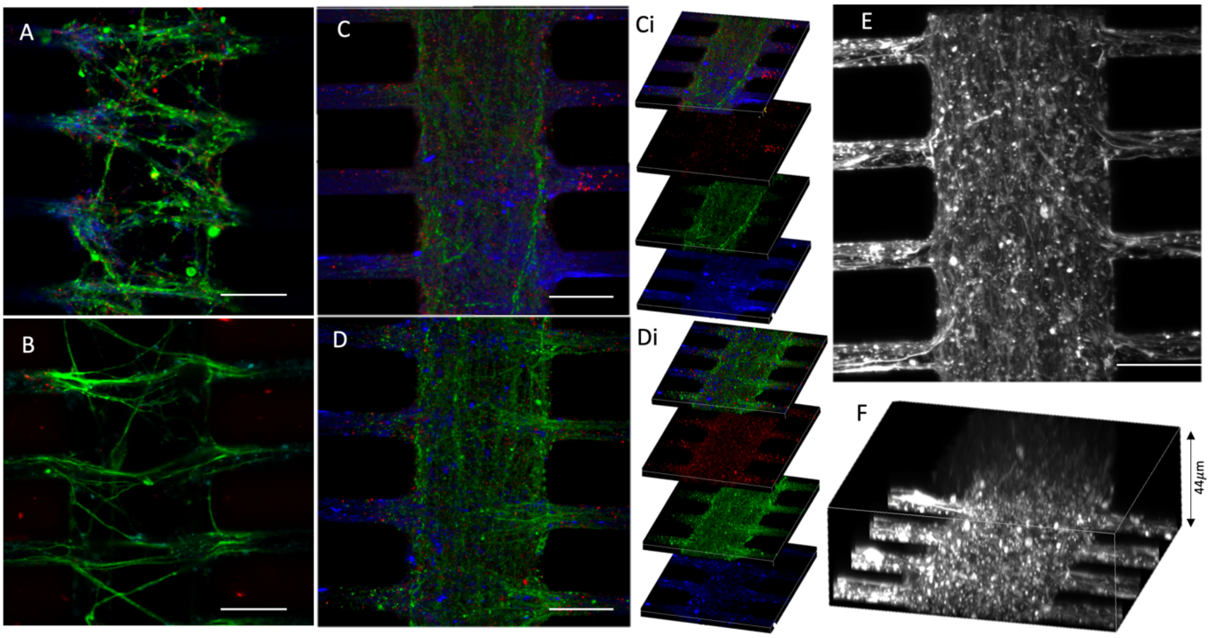


**Fig.S2 Difference in neuritic density**. **A** and **B**) Example images from the control neural networks fluorescently labelled with the presynaptic marker Piccolo (green), the post synaptic marker PSD95 (red) and the f-actin filament marker Phalloidin (blue) (30µm scale bar). **C** and **D**) Equivalent example images from the LRRK2 neural networks. **Ci** and **Di**) show the 10µm thick z-stacks compiled into **C** and **D**, respectively. **E** and **F** show an LRRK2 neural network labelled with phalloidin, where a z-stack has been taken to capture the entire volume of neurites contained within this section of the synaptic compartment, where **F** shows the volumetric sideview illustrating a neuritic bundle thickness of 44µm.

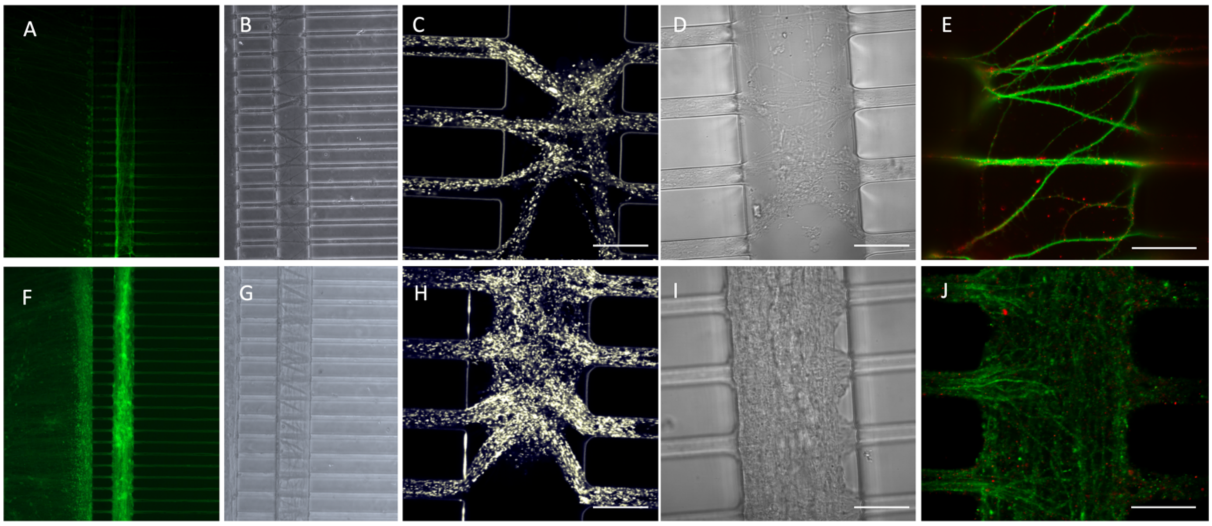


**Fig.S3 Difference in neural network morphology visible at all levels. A-E)** Example images from different control neural networks during each assay, while **F-J**) Equivalent images from the LRRK2 neural networks. There is a striking difference in the neurite outgrowth profile within the synaptic compartments, with clear fasciculation observed in **E)** control networks as opposed to **J)** aberrant growth in LRKK2 ones. All images have a 30µm scale bar, apart from E and J, which have a 20mµ scale bar.

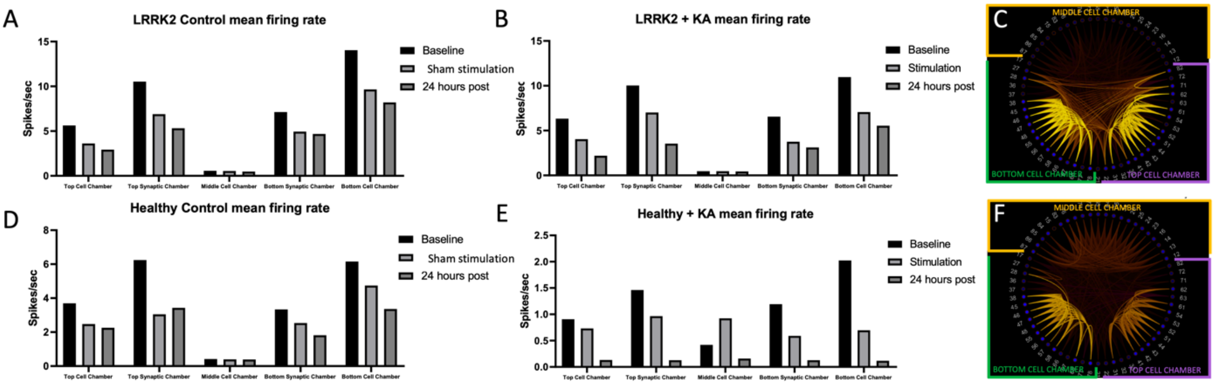


**Fig.S4 Reproducibility of network activity traits and functional connectivity through structuring using a multi-nodal microfluidic chip.** Mean firing rate (MFR) bar graphs organized by the different microfluidic chip areas, of both the (**A**) LRRK2 and (**D**) healthy neural network control groups clearly demonstrate that the activity of the structured neural networks is reproducibly influenced by the microfluidic chip physical structuring. Furthermore, the same trend is visible at the baseline timepoint of both the (**B**) LRRK2 and (**E**) healthy neural networks receiving KA. Although the MFR of the LRRK2 neural networks is far greater than the MFR of the healthy neural networks (relative x-axis), they follow the same trend, with much greater network activity measured at the top and bottom cell chambers than in the middle cell chamber. This is also visible from the correlation scheme balls presented in **C** and **F** of a baseline measurement from the LRRK2 and healthy neural networks, respectively, where greater correlations are found between the activity measured within each chamber than between, and stronger correlation found in the top and bottom cell chamber than in the middle.

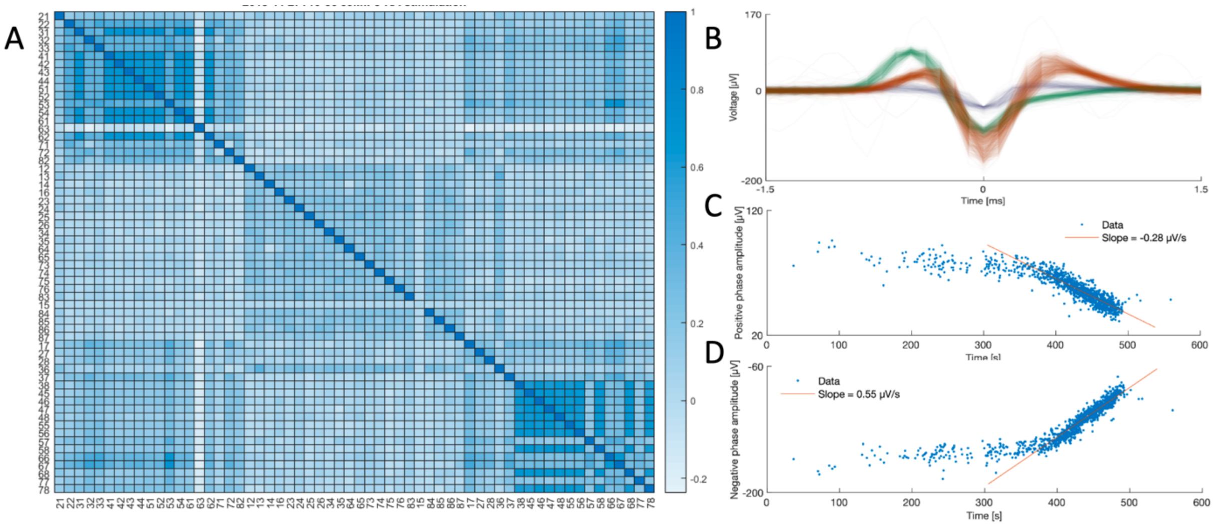


**Fig.S5 Electrophysiological activity in LRRK2 neural network during KA stimulation. A)** shows the correlation map of the activity measured during KA stimulation of the LRRK2 neural network shown in **Fig.6E-G**. **B**) shows the three distinct spike clusters measured at a randomly chosen electrode (number 63) of this network during the stimulation period. **C** and **D**) respectively show the trailing positive and negative phase amplitudes of the spikes from the neuron represented in red in **B** over time. The firing rate of this neuron can be seen to increase drastically 300-500 s into the recording session (see also **Fig.6C**), firing at the maximal rate allowed by the refractory period. This period coincides with both the negative and positive amplitudes of its signal becoming progressively reduced, followed by an abrupt silencing. This electrode is located in the top, stimulated cell chamber. Interestingly, from the MFR map in **A** it is clear that it is the only electrode that is negatively correlated (mean r = -0.2) with the activity of some other electrodes located in the same chamber, as well as some electrodes in the bottom cell chamber, indicating that the neuron whose spikes are plotted in red in **B** exerts an inhibitory effect.

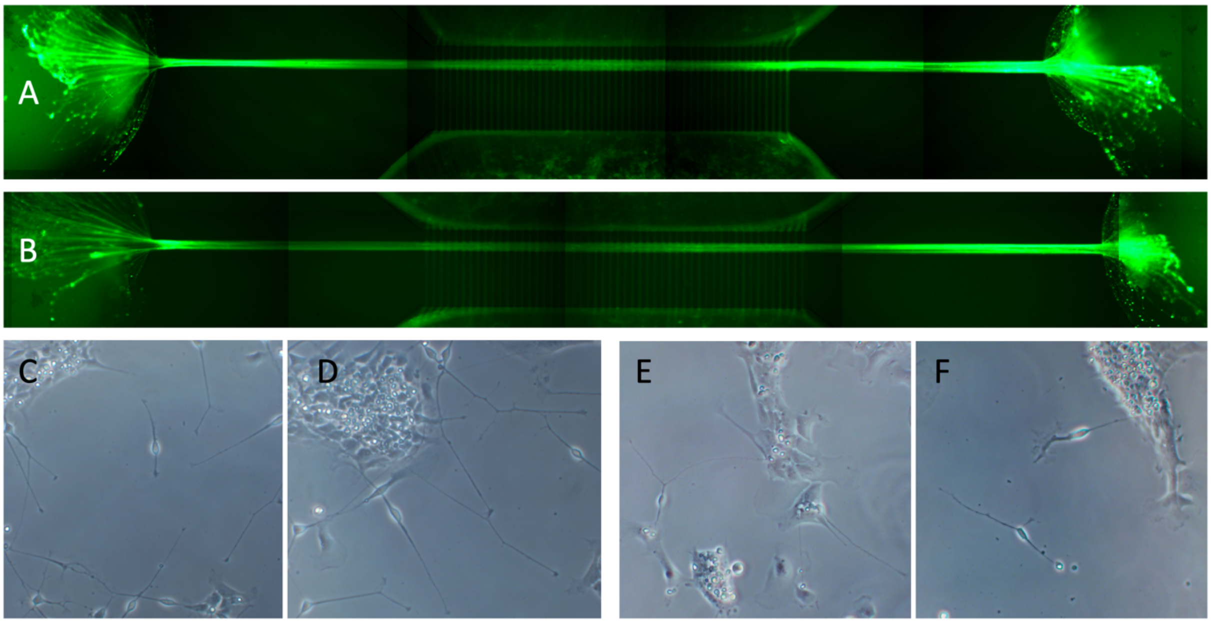


**Fig.S6**. **Aberrant neurite outgrowth and growth cone differences in LRRK2 neural networks versus controls. A,B**) show LRRK2 neural networks labelled with the fluorescent Calcein-AM, where massive bundles of neurites can be seen exiting at both ends of the synaptic compartments (the inlet and outlet), where there are no cells to connect up with. These bundles are perpendicular to the axon tunnels interconnecting the cell chambers. **C,D**) shows brightfield images of LRRK2 NSCs, and **E,F**) of control neural networks after 1 day of differentiation, demonstrating differences in neurite outgrowth and growth cone profile, i.e. qualitatively more growth cones and branching neurites present in the LRRK2 neurons compared to controls.

### Supplementary information on cell lines and culturing (Materials and Methods)

iPSC-derived H9N neural stem cells (ax0019) (control) and iPSC-derived H9N neural stem cells homozygous inserted with the LRRK2 G2019S (ax0310) purchased form Axol Bioscience were used for the experiments. Both lines are derived from the same donor dermal fibroblasts (female, 64 yr). General gene expression profiles from Axol's iPSC-derived NSCs is published in Gene expression omnibus (GEO) of the National Center for Biotechnology information (NCBI) (https://www.ncbi.nlm.nih.gov/geo/query/acc.cgi?acc=GSE61358 ).

Furthermore, gene editing and genotyped example data, as well as the CRISPR plasmid donor scheme for the LRRK2 line (ax0310) is shown below with the permission of Axol Bioscience.

| Project | # Clones | # KI/KI | # KI/+ | # KI/- | other |
|---|---|---|---|---|---|
| LARRK2 | 31 | 2 | 6 | 4 | 16 |

- 1 (KI/KI) and 2 (KI/+) clones fully genotyped; example data below:

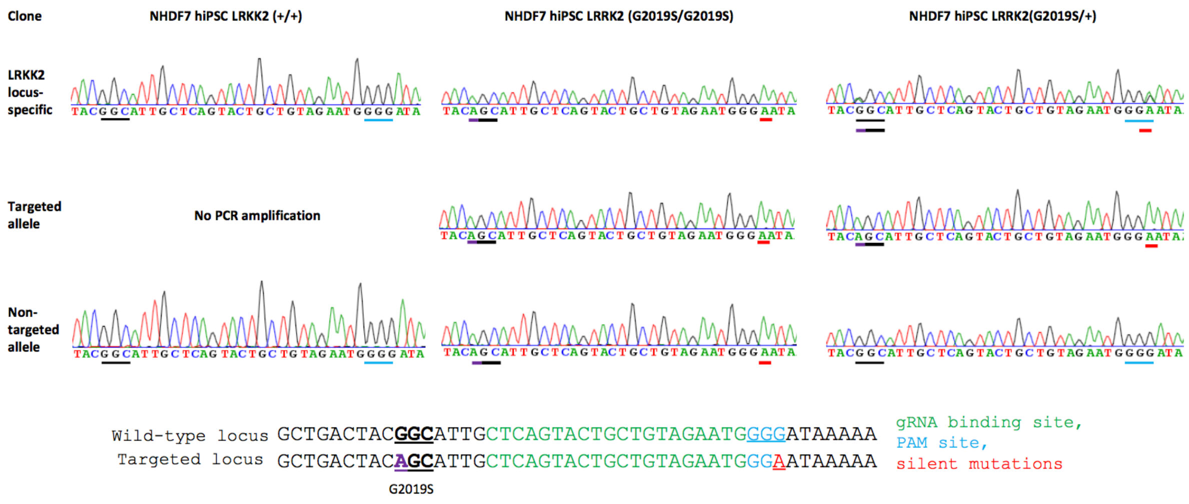

### LRRK2 G2019S

➢ **CRISPR + plasmid donor**

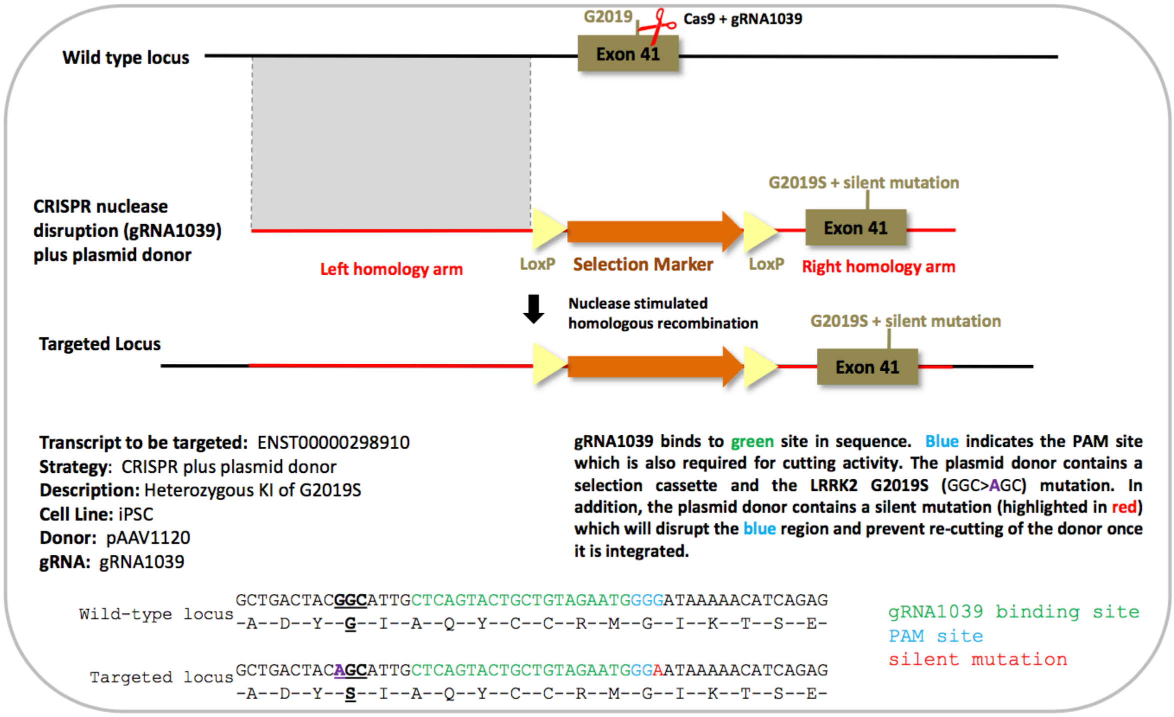


**Protocol for culture of H9N NSCs (ax0019 and ax0310), adapted from Axol Bioscience's protocol (Human iPSC-derived Neural Stem Cell master protocol (version 5.0)**
**System A**

<u>List of media and supplements</u>
Neural Expansion -XF Medium (ax0030-500)
Neural Maintenance - XF Medium (ax0032-500)
Neual Differentiation – XF Medium (ax0034-125)
Sure Growth recombinant human FGF2 (ax0047)
Sure Growth X recombinant human EGF (ax0047X)
0.01% Poly-L-Ornithine solution (PLO) (Sigma, P4967)
Natural Mouse Laminin (Thermo Fisher Scientific, 23017015)
L15 Lebovitz medium (Sigma, L5521)
KnockOut serum replacement (KOSR, Thermo Fisher Scientific, 10828010)

<u>L15/laminin coating solution</u>
3ml L15 Medium
48ul Natural mouse laminin (1mg/ml)
75ul Sodium Bicarbonate

<u>Day before seeding:</u>
Coat one well of a 6-well plate with PLO for 2 hours in incubator, wash with sterile MQ water, and leave overnight with L15-laminin in fridge.
Thaw an aliquot of Expansion medium (from -80 freezer) overnight in fridge
(Protect from light, can be kept for 1 week if supplemented (EGF, FGF2), 2 weeks if not)

Day of seed:
Prepare Neural Expansion medium with supplements (for spinning and seeding).

10ml Neural Expansion medium
2ul FGF2 (20ng/ml) (stock is 100ug/ml)
2ul EGF (20ng/ml)
10ul Rock Inhibitor (only when thawing/ splitting)
100ul Pen-Strep

Aspirate coating and add Neural Expansion medium to the well (ca 1,5ml) and return to incubator until seeding.
Thaw vial of cells in water bath (no shaking)
Pre-coat pipette with KOSR, (dropwise add cells to tube containing 10ml warm expansion medium. Spin for 200g x 5 min. Aspirate supernatant, precoat pipette with KOSR, resuspend cells in 0,5-1ml expansion medium and seed in 6-well.

Expansion:
Every 2 days, replace all the medium with Neural Expansion medium (supplemented with FGF and EGF). When the culture is 70-80% confluent, they are ready to undergo passage. This usually takes a long time (> one week). Do not expand more than 3 passages.

Splitting:
1/2 split. Coat 2 6-well wells.
Thaw a vial of Axols "Unlock" (aliquoted and stored in -80 in a box at the bottom).
Prepare Expansion medium (+ Rock inhibitor).
Aspirate media and rinse surface with DPBS--. Add Unlock (1ml /10cm^2), place in incubator for 5 min. Use 4x (unlock volume) of Neural expansion medium to stop reaction. Spin for 200g x 5min. Aspirate and resuspend in supplemented Neural Expansion medium, seed 50% in each of the 6-wells.

Synchronous Differentiation
When you have expanded long enough and are ready to start differentiating:
Full media change to Neural Expansion medium without supplements. Thaw aliquot of Neural Differentiation medium overnight in fridge.
24 hours later, conduct a full media change to Neural Differentiation medium (no supplements)
Every three days change 50% of the medium with fresh Neural Differentiation medium. A pure neuronal population takes anywhere between 3-10 days to achieve (dependent on the confluency, <60% = 3 days ++, >60% confluency takes up to 10 days, fully confluent may never become pure).

Maintenance
When fully differentiated, thaw an aliqot of Neural Maintenance medium overnight in fridge. Replace half of the medium with Neural Maintenance medium. 24 hours later, replace half the medium with Neural Maintenance.
After this, replace 50% of the medium with Neural Maintenance medium every 3 days. Might have to add extra laminin if the cells start detaching.
